# Dissecting the homeodomain *MAT* locus and engineering novel tripolar and bipolar mating systems in *Cryptococcus amylolentus*

**DOI:** 10.1101/2025.11.14.688536

**Authors:** Liping Xiong, Vikas Yadav, Sheng Sun, Joseph Heitman

**Author notes:** Corresponding author: Joseph Heitman.

## Abstract

Sex in fungi is governed by the mating-type (*MAT*) locus, which exists as bipolar, pseudobipolar, or tetrapolar systems. The significance and impact of *MAT* on sexual reproduction, however, remain understudied. Furthermore, the evolution of fungal *MAT* loci shares features with the evolution of sex chromosomes in plants and animals. Pathogenic *Cryptococcus* species harbor a bipolar system with a large contiguous *MAT* locus, whereas closely related species, such as the non-pathogen *C. amylolentus* possess a tetrapolar system with unlinked *P/R* and *HD* loci. The *HD* locus encodes homeobox domain containing proteins that play an important and evolutionarily conserved role in sexual reproduction. Here, we explored the roles of *HD* genes in sexual reproduction and determined the implications of a tetrapolar to bipolar *MAT* transition. With a CRISPR–Cas9 system we developed for *C. amylolentus*, we generated gene deletion mutants and demonstrated that a single compatible Sxi1-Sxi2 pair is necessary and sufficient for mating. By relocating the *HD* genes to the *P/R* locus, we found that the artificially generated bipolar configuration led to defective sexual development, which could be partially restored through additional rounds of sexual reproduction. Transcriptomic profiling further revealed that a Sxi1-Sxi2 heterodimeric complex drives expression of genes required for DNA replication and ergosterol biosynthesis during sexual reproduction. These findings provide the first experimental demonstration of a tetrapolar-to-bipolar transition in a tetrapolar mating species, illuminating *MAT* locus evolution and homeodomain protein functions in *Cryptococcus*.

**Importance:** Sexual reproduction is critical for fungal survival and adaptation, yet the mechanisms driving transitions between mating systems remain unclear. With *Cryptococcus amylolentus*, we provide the first experimental validation of a mating system transition from its original tetrapolar to an intermediate tripolar to a derived bipolar in a tetrapolar species. We show that HD heterodimers phenotypically govern dikaryotic filamentation and also transcriptionally modulate DNA replication. These findings establish a mechanistic basis for how *MAT* locus reorganization drives bipolar evolution from an ancestral tetrapolar state and reinforce that fertility depends on the coordinated control of *MAT* locus architecture and regulatory functions.

## Introduction

Sexual reproduction in fungi plays a pivotal role in promoting genetic diversity, maintaining genome integrity, and facilitating adaptation to dynamic environments. This process is governed by the mating-type (*MAT*) locus, a specialized genomic region that encodes key determinants of mating compatibility and sexual development (1–3). *MAT* loci are diverse in length, ranging from a few kilobases to a few megabases and even spanning an entire chromosome. *MAT* loci are also diverse in gene content and structure, even among closely related species (4). The organization of *MAT* loci is the fundamental determinant of fungal mating systems. Fungi typically exhibit one of three mating system configurations—bipolar, pseudobipolar, or tetrapolar—with all three commonly found among basidiomycetes. Most basidiomycete species are tetrapolar in which the mating types are determined by alleles at two unlinked loci: a P/R locus (aka. A locus) encoding pheromones and pheromone receptors mediating partner recognition and cell–cell fusion, and an HD locus (aka. B locus) encoding transcription factors controlling cell type identity and post-fusion sexual development (5). Successful mating in tetrapolar species, such as the corn smut fungus *Ustilago maydis* (6), the mushroom-forming *Coprinopsis cinerea* (7), and *Schizophyllum commune* (8), requires partners to differ at both loci, resulting in potentially up to thousands of distinct mating types and presumably promoting high outcrossing efficiency. In contrast, in bipolar species, such as the sugarcane smut fungus *Sporisorium scitamineum* (9) and the human fungal pathogen *Cryptococcus* species complex (1, 3), the two loci are physically linked into a single *MAT* locus, potentially increasing the likelihood of inbreeding.

Interestingly, transitions between tetrapolar, bipolar, and intermediate pseudo-bipolar mating systems appear to have occurred repeatedly and independently across diverse fungal lineages and within closely related species (5, 10–15). For example, the tetrapolar fungus *U. maydis* carries unlinked *P/R* and *HD* loci on separate chromosomes, whereas its sister species *Ustilago hordei* and *Ustilago bromivora* have evolved bipolar systems with the two *MAT* loci located on the same chromosome, separated by ∼500 kb and ∼180 kb, respectively (16, 17). Similarly, several human skin fungal pathogens in the *Malassezia* species complex (*M. globosa*, *M. furfur*, and *M. yamatoensis*) exhibit bipolar systems with the two *MAT* loci separated by variable distances ranging from ∼140 kb to ∼580 kb (18, 19). Notably, *M. sympodialis* represents a pseudo-bipolar state, in which the *P/R* and *HD* loci are physically linked yet ∼141 kb apart and still undergo recombination (14, 15, 19). The coexistence of these distinct strategies, often even within closely related taxa, highlights the remarkable plasticity of fungal mating systems and suggests that transitions occur frequently, likely mediated by chromosomal rearrangements and recombination suppression at *MAT* loci.

Among basidiomycete fungi, *C. neoformans* is a well-established model for investigating both development and pathogenesis. Its sexual cycle can be readily analyzed with robust genetic tools, and its clinical significance is underscored by its role as a leading cause of meningoencephalitis in immunocompromised patients. *Cryptococcal* meningitis affects ∼194,000 people with ∼147,000 deaths (∼76% mortality) globally each year (20, 21). The primary infectious propagules of *Cryptococcus* species are desiccated yeast cells and spores, the latter of which are produced during sexual reproduction. Species in the pathogenic *Cryptococcus* species complex all exhibit a bipolar mating system, with a large bi-allelic *MAT* locus (>120 kb, *MAT***a** and *MAT*α) that evolved from fusion of the ancestral *P/R* and *HD* loci (15, 22, 23). In *C. neoformans* the *MAT* locus is also associated with virulence, with *MAT*α consistently associated with greater pathogenicity in both clinical and animal studies (24, 25).

In contrast, the closely related non-pathogenic species *C. amylolentus* retains the ancestral tetrapolar system, with a *P/R* locus (∼96 kb) on chromosome 10 and an *HD* locus (∼22 kb) on chromosome 11 that act independently (26–28). The *HD* locus of *C. amylolentus* contains two divergently oriented *HD* genes, *SXI1* (i.e. *HD1*) and *SXI2* (i.e. *HD2*). Sexual reproduction occurs through heterothallism, requiring mating partners with compatible *MAT* alleles at both the *P/R* and *HD* loci (26, 29). It is hypothesized that the Sxi1 and Sxi2 proteins from the mating partners form heterodimers in the zygote, but it is not yet clear whether both possible heteroallelic heterodimers in the zygote are required for successful sexual development, or whether homoallelic heterodimers could also be formed between Sxi1 and Sxi2 of the same strain. Interestingly, many genes within the fused *MAT* locus of pathogenic species remain syntenic with those near the *MAT* loci of *C. amylolentus* (26). This close relationship suggests that shifts between tetrapolar and bipolar systems may be driven by chromosomal rearrangements and selection for reproductive efficiency (5, 28, 30).

It has been hypothesized that the fused *MAT* locus in pathogenic *Cryptococcus* species arose from ancestral *P/R* and *HD* loci through ectopic recombination (15, 28). Experimental relocation of the *SXI1*α and *SXI2***a** *HD* genes in *C. neoformans* demonstrated that engineered strains can complete a tetrapolar sexual cycle (12). However, no experimental validation has as yet been carried out in a naturally tetrapolar species. More recently, chromosomal translocation events involving those carrying the *MAT* loci were identified, establishing the initial linkage between *P/R* and *HD* loci in the pathogenic *Cryptococcus* species complex (15, 28). As a closely related tetrapolar species, *C. amylolentus* thus provides a compelling model in which engineering a synthetic shift from tetrapolar to bipolar organization enables direct tests of this evolutionary scenario observed in pathogenic lineages.

Another related major unresolved question concerns the molecular mechanisms of *HD1*-*HD2* function and their downstream targets. Many basidiomycetes encode multiple *HD1*-*HD2* pairs that differ in strength or specificity (8, 31), yet across species these heterodimers consistently act as master regulators, triggering transcriptional cascades that govern mating, filamentation, and sexual differentiation (31–34). For example, pathogenic *Cryptococcus* species have a single *HD1*-*HD2* pair in the zygote composed of Sxi1α and Sxi2**a**, which is necessary and sufficient for inducing hyphal development, a key step in sexual development (33–35). Thus far, only a few downstream genes have been identified (36, 37). For example, in *Ustilago maydis*, the bE/bW heterodimer directly regulates *RBF1* encoding a key sexual development factor (13). However, a homolog of *RBF1* in *Cryptococcus neoformans*, *ZNF2* (a zinc-finger transcription factor gene), is required for filamentation but is not a target of the HD transcription factor complex (38). Instead, the *CLP1* (required for dikaryotic filament formation) (39, 40), *CPR2* (a constitutively active G protein coupled receptor gene) (41), and *VAD1* (encodes a RNA binding protein) genes (20, 42) are directly regulated by the Sxi1α-Sxi2**a** complex, highlighting the species-specific nature of *HD1*–*HD2* regulons. These differences underscore the need to investigate HD proteins across diverse fungi, as such studies may reveal conserved core developmental pathways controlled by HD heterodimers.

In this study, we investigated the functional consequences of the *HD* genes (*SXI1* and *SXI2*), as well as the impact of their physical location, on sexual development of the tetrapolar species *C. amylolentus*. With our newly developed CRISPR-Cas9 system, we first generated strains with deletions of *SXI1*, *SXI2*, or both in the two mating compatible strains CBS6039 and CBS6273. While deleting the *HD* genes did not affect vegetative growth or virulence traits, mutants lacking both *HD* genes were defective in mating, suggesting that at least one heteroallelic Sxi1-Sxi2 pair is essential for sexual reproduction. Our RNA-seq analysis suggests that Sxi1-Sxi2 heterodimers may coordinate S-phase entry and activate the replication machinery and lipid biosynthesis in preparation for subsequent sexual development. We then relocated both *HD* genes from the *HD* locus into the *P/R* locus, and generated *P/R*+*HD* strains (i.e. *MAT*-fused). We showed that these *P/R*+*HD* strains can undergo successful sexual reproduction with wild-type tetrapolar strains possessing compatible *MAT* alleles in “tripolar” matings, producing viable basidiospores. However, when the engineered *P/R+HD* strains with compatible *MAT* alleles were crossed with each other in “bipolar” matings, we observed reduced cell-cell fusion and defective hyphal growth and basidiospore production, implying phenotypic effects on sexual reproduction by the endogenous genomic environment of the *HD* locus.

Taken together, our findings provide the first experimental demonstration of a complete reconfiguration of the *MAT* loci, from the original tetrapolar system to tripolar, and then bipolar, revealing the functional implications of the *HD* locus during sexual reproduction, with insights on the organization and evolution of *MAT* loci in basidiomycetes.

## Results

### One pair of heteroallelic Sxi1-Sxi2 homeodomain factors is necessary and sufficient for successful mating

In *C. neoformans*, Sxi1α and Sxi2**a** form an obligate heterodimer after mating, which acts as a master regulator of the dikaryotic phase, essential for driving hyphal development, karyogamy, and meiosis (33). In *C. amylolentus*, because the *HD* locus contains both *HD1* and *HD2*, there are three different scenarios as to how the sexual development of the zygote could be regulated by the *HD* heterodimers. HD dimers could be formed by: 1) the HD proteins from the same parent (i.e. homoallelic dimer); 2) HD proteins from the two parents (i.e. heteroallelic dimer); or 3) the collective actions of two heteroallelic dimers in the zygote.

To test these hypotheses, we first developed and optimized a CRISPR-Cas9 system for *C. amylolentus* for genetic manipulation, with which we then constructed strains with single (*sxi1*Δ and *sxi2*Δ) and double (*sxi1*Δ *sxi2*Δ; hereafter referred to as *hd*Δ for simplicity) deletions of the *HD* genes in both the CBS6039 (with *NAT* dominant marker) and CBS6273 (with *NEO* dominant marker) backgrounds (Figure S2). We then set up these *HD*-deletion strains, together with the wild-type background strains, in all possible pair-wise crosses (Figure 1A) between strains of opposite mating types. Under mating conditions, solo cultures of *C. amylolentus* wild type isolates produced robust vegetative hyphae, while co-cultures of strains with compatible mating types produced a mixture of both vegetative hyphae and sectors producing sexual/dikaryotic hyphae, basidia, and spores (Figure 1A and 1B).

**Figure 1.**
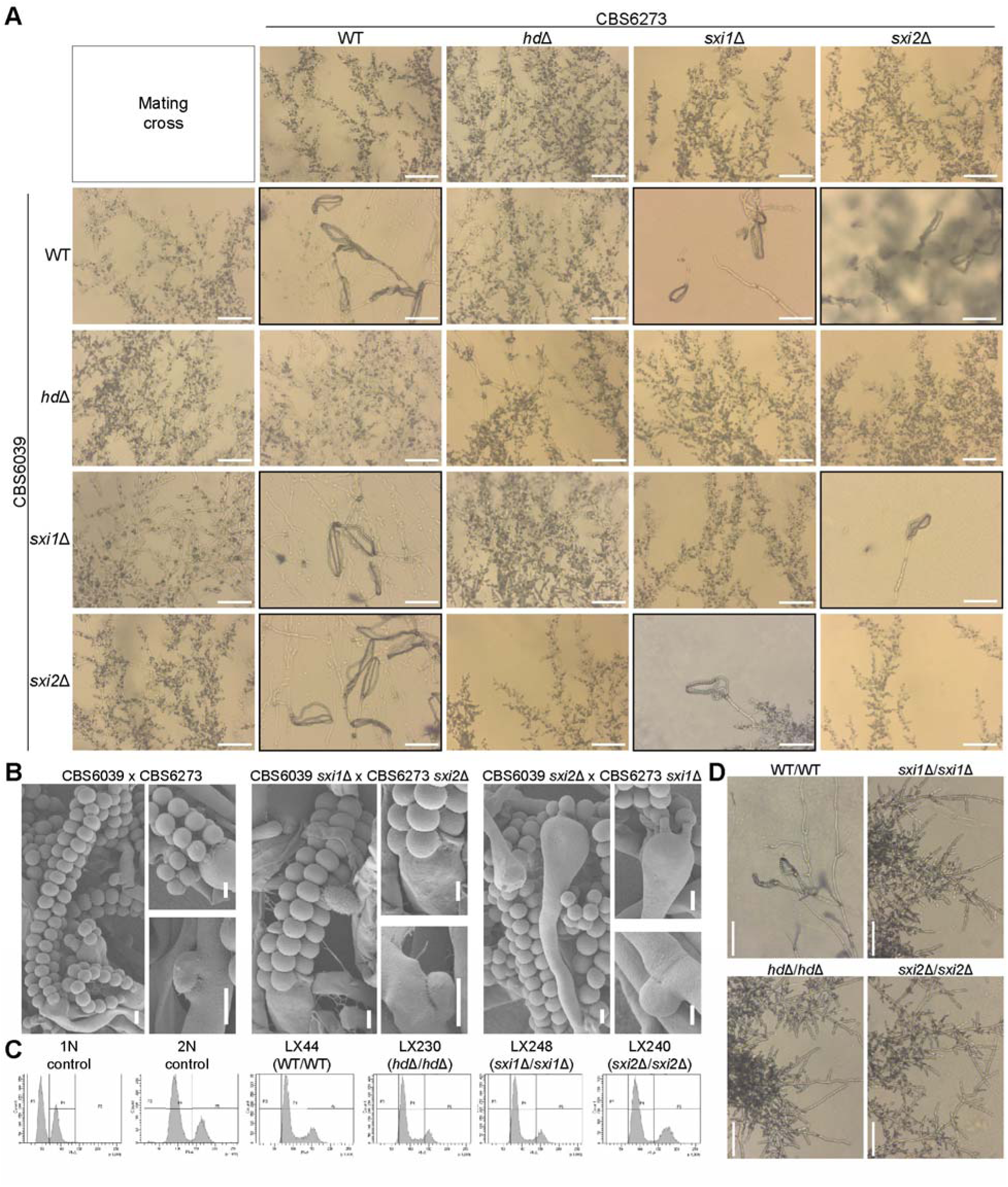
One pair of heteroallelic HD1-HD2 is required and sufficient for mating. **(A)** Mating phenotypes of bilateral crosses between *C. amylolentus* wild type strains and their *HD* gene deletion mutants. Microscopic examination of mating structures produced during a cross between the indicated strains on V8 (pH=5) medium incubated in the dark at room temperature for 2 weeks. Scale bar: 50 µm. Black boxes highlighted basidia spores that were observed; red boxes highlight defective mating phenotypes. **(B)** SEM examination of mating structures produced during mating between the two *C. amylolentus* wild type strains, CBS6039 and CBS6273, and their *sxi1*Δ and *sxi2*Δ mutants on V8 (pH=5) medium incubated in the dark at room temperature for 2 weeks. Upper panel: basidia and long spore chains. Bottom panel: magnified images of basidiospores attached to basidia and clamp connections. Scale bar: 2 µm. **(C)** FACS analysis. Haploid H99 served as a haploid control, and CnLC6683 as a diploid control. LX44 is a diploid isolate from LX21 (CBS6039 Nat^R^) x LX26 (CBS6273 Neo^R^), LX230 is a diploid isolate from LX68 (CBS6039 *hd*Δ) x LX148 (CBS6273 *hd*Δ), LX248 is a diploid isolate from LX78 (CBS6039 *sxi1*Δ) x LX153 (CBS6273 *sxi1*Δ), and LX240 is a diploid isolate from LX74 (CBS6039 *sxi2*Δ) x LX135 (CBS6273 *sxi2*Δ). **(D)** Microscopic examination of mating structures of the indicated diploids on V8 (pH=5) medium incubated in the dark at room temperature for 1 week. Clamp cells, basidia, and spores were observed in LX44 (WT/WT). Scale bar: 50 µm.

Neither of the two *hd*Δ strains was able to mate with any of the strains of the opposite mating type, including both wild-type and the *HD* deletion strains (Figure 1A), showing that successful mating requires *HD* genes from both mating types in the zygote. For strains with deletions of a single *HD* gene, they were able to undergo successful mating with all of the strains of the opposite mating type, except those containing allelic deletions of the *SXI1* or *SXI2* genes (Figure 1A), further demonstrating that heterodimers formed by the *HD* genes from the two parental strains (i.e. heteroallelic dimers) are required for successful mating, and only one such heterodimer is sufficient. Additionally, we did not observe any difference in mating properties between the deletion strains of the two *HD* genes (i.e. *sxi1*Δ vs. *sxi2*Δ), suggesting there is no preference between the two possible heteroallelic *HD*-dimers.

Failed mating could be due to defects in cell-cell fusion or sexual development after zygote formation. To investigate the nature of the failed sexual reproduction that was observed in the *HD*-deletion strains, we constructed diploid homozygous deletion strains lacking *SXI1*, *SXI2*, or both, by fusing the CBS6039 and CBS6273 strains containing the *HD* gene deletions (*sxi1*Δ/*sxi1*Δ, *sxi2*Δ/*sxi2*Δ, or *hd*Δ/*hd*Δ) (Figure 1C). None of these strains was able to undergo sexual development under mating inducing conditions (Figure 1D). The fact that strains with deletions of these genes could fuse and form diploids, but the resulting zygote failed to undergo further sexual development, suggests that the mating defect caused by the deletion of these *HD* genes is not during cell-cell fusion, but is instead post-fusion because Sxi1-Sxi2 is required for the zygote to undergo sexual development.

Why does successful mating require heteroallelic HD dimers? One possibility could be that the HD1 and HD2 proteins only form heterodimers between alleles of the opposite mating types. To test this, we employed AlphaFold modeling to predict potential interactions among the four HD proteins. No homodimer formation was predicted for the Sxi1 or Sxi2 proteins. In contrast, both heteroallelic and homoallelic Sxi1-Sxi2 dimers were predicted, with the heteroallelic dimers showing higher confidence scores than the homoallelic dimers (Figure S1). The high sequence similarity (∼92%) between Sxi1 proteins, as well as between Sxi2 proteins, from the two parental isolates may account for the predicted homoallelic dimers. Closer comparisons revealed that homoallelic dimers differed markedly in structure, whereas the heteroallelic dimers are structurally similar (Figure S1). Consistent with the non-selfing nature of each parental isolate, these results suggested that homoallelic dimers, even if formed, are insufficient to drive the sexual development in *C. amylolentus*.

Taken together, our results demonstrate that one of the two possible heteroallelic Sxi1-Sxi2 dimers is necessary and sufficient for successful sexual development after cell-cell fusion in *C. amylolentus*.

### Relocation of the *HD* locus did not affect mating in a tripolar *MAT* configuration

In the wild-type *C. amylolentus* strains CBS6039 and CBS6273, the *P/R* (A) and *HD* (B) *MAT* loci are located on chromosomes 10 and 11, respectively. We studied how changing this tetrapolar configuration could affect sexual reproduction. To this end, we first identified “safe haven” (SH) intergenic regions in the *P/R* locus of the two strains that lack any repeats, transposons, or DNA/RNA/protein-binding domains based on NCBI’s Conserved Domain Database. The SH sites are different between the two strains due to the highly rearranged nature of the *P/R MAT* alleles, and are located between the *RPL39* gene and the 5’ edge of the *P/R* in CBS6039 and between the *CID1* and *GEF1* genes in CBS6273 (Figure 2A). We then inserted the *SXI1-SXI2* gene pair into the SH sites in the *hd*Δ strains that we previously constructed in the CBS6039 and CBS6273 backgrounds, generating the *P/R*+*HD* (i.e. A+B or *MAT*-fused) strains. To control for introducing exogenous sequences into the SH sites, we also inserted dominant selectable markers into the SH sites (*SH*-marker) of the CBS6039 and CBS6273 wild-type strains.

**Figure 2.**
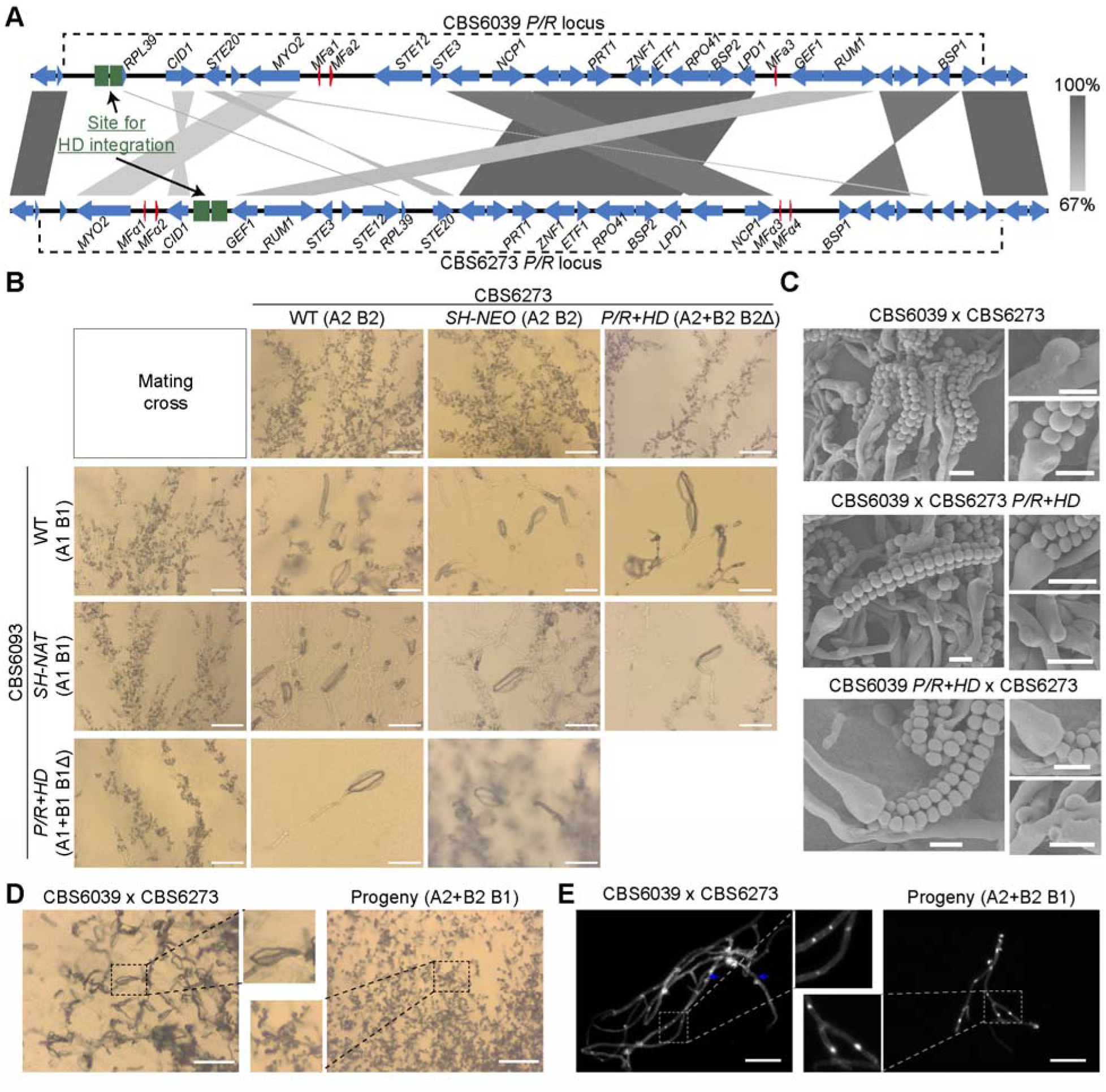
Relocation of the *HD* locus enables mating in a tripolar *MAT* configuration in *C. amylolentus*. **(A)** Schematic graph of the *HD* relocation to *P/R* loci. Shown here are results of synteny analyses between genomic sequences from strains CBS6039 and CBS6273 for the *P/R* loci based on nanopore sequencing. Red color highlights the genes that define the *P/R* locus (mating pheromones and *STE3*). *HD* was relocated into *SH* regions that are inside the boundary of the *P/R* locus in both CBS6039 (between the 5’ edge of *P/R* and *RPL39* gene) and CBS6273 (between the *CID1* and *GEF1 gene*), generating *P/R+HD* fused (bipolar) isolates in both strain backgrounds. Green boxes indicate the homologous flanking regions employed for strain construction. **(B)** Mating phenotypes of all possible strain combinations between *C. amylolentus* wild type strains and their *P/R+HD* mutants. All of the solo and cocultures were grown on V8 (pH=5) medium incubated in the dark at room temperature for 2 weeks. Scale bar: 50 µm. Black boxes highlight basidia spores that were observed between indicated mating crosses. **(C)** SEM revealed basidiospores attached to basidia and unfused clamp connections. Samples prepared for SEM were collected from those of (B). Scale bar: 2 µm. **(D)** Progeny with an active Sxi1-Sxi2 heterodimer are self-filamentous. Shown here are the phenotypes of a wild-type cross (A1 B1 x A2 B2) and a progeny carrying both B1 and B2. Mating assays were set up on V8 medium (pH=5) for 2 weeks of incubation. Scale bar: 100 µm. (**E)** Hyphae after DAPI (4,6-diamidino-2-phenylindole) staining. Hyphal cells were harvested from the (D). Filaments formed by the progeny (A2+B2 B1) are monokaryons (right), in contrast to dikaryons observed in the sexual hyphae formed by the wild-type cross (left). White arrows indicate the nuclei stained white; blue arrows indicate the clamp cells. Scale bar: 100 µm.

We next analyzed the mating properties of these *MAT*-fused strains, by conducting all possible pair-wise mating assays between strains in the CBS6039 and CBS6273 backgrounds that are of the wild-type, *hd*Δ, *SH*-marker, and *P/R*+*HD* genotypes at the *MAT* loci (Figure 2B). The *SH*-marker strains showed the same mating property as their corresponding wild-type background strains, indicating that the insertion of exogenous sequences into the *SH* loci did not disrupt the functionality of the *P/R* alleles (Figure 2B). Consistent with previous results, the *hd*Δ strains in both backgrounds failed to mate with any strain, reaffirming that successful mating requires the *HD* genes from both mating partners. Interestingly, the *MAT*-fused strains (*P/R*+*HD*) (in which the *HD* genes were reintroduced into the *P/R* locus of the sterile *hd*Δ strains) were now able to mate with the strains of the opposite mating type, and produced sexual development structures, including sexual hyphae, clamp cells, basidia, and basidiospores, similar to the wild-type stains (Figure 2B and 2C). Thus, the *MAT*-fused strains were able to successfully mate with the strains in which the *P/R* and *HD* are unlinked, constituting a “tripolar” mating.

We next recovered basidiospores from individual basidia of four reciprocal crosses between wild-type strains and the *MAT*-fused strains (Table 1). Specifically, we dissected a total of 20 basidia (10 basidia each) from two crosses between CBS6039 *MAT*-fused strains (LX143 and LX283, genotype “A1+B1 B1Δ”) and the CBS6273 wild-type strain (A2 B2). With the exception of two basidia (No. 8 in LX143 cross and No. 2 in LX283 cross), the vast majority of basidia produced spores with germination rates higher than 20%, which is comparable to previous studies (28, 43). Analyses of the mating types showed that all four possible mating types (A1+B1 B1Δ, A2 B2, A1+B1 B2, and A2 B1Δ) were present among the basidiospores, with the first two being the parental types and the last two being the recombinants (Table 1). We also dissected a total of 19 basidia from two reciprocal crosses between CBS6039 wild-type strain (A1 B1) and CBS6273 *MAT*-fused strains (LX167 and LX170, genotype “A2+B2 B2Δ”). We observed similar spore germination rates, and again, we identified all four mating types among the basidiospores, suggesting segregation of the *MAT* alleles in these “tripolar” matings.

**Table 1.** Genotyping analysis of spores dissected from tripolar crosses.

| Cross | Basidia <sup>1</sup> | Dissected<br>spores | Germinated<br>spores | Germination<br>rate (%) | Progeny genotype |  |  |  |
| --- | --- | --- | --- | --- | --- | --- | --- | --- |
|  |  |  |  |  | A1+B1 B1Δ | A2 B1Δ | A1+B1 B2 | A2 B2 |
| LX143 (A1+B1 B1Δ) |  |  |  |  |  |  |  |  |
| □ |  |  |  |  |  |  |  |  |
| CBS6273 (A2 B2) |  |  |  |  |  |  |  |  |
|  | 1 | 7 | 2 | 29 | 0 | 1 | 1 | 0 |
|  | 2 | 10 | 2 | 20 | 0 | 1 | 1 | 0 |
|  | 3 | 14 | 13 | 93 | 5 | 6 | 1 | 1 |
|  | 4 | 11 | 7 | 64 | 2 | 0 | 2 | 3 |
|  | 5 | 10 | 6 | 60 | 0 | 2 | 2 | 2 |
|  | 6*** | 14 | 12 | 86 | 2 | 3 | 4 | 3 |
|  | 7*** | 12 | 12 | 100 | 3 | 4 | 3 | 2 |
|  | 8 | 10 | 1 | 10 | 0 | 0 | 0 | 1 |
|  | 9 | 8 | 2 | 25 | 0 | 1 | 1 | 0 |
|  | 10 | 10 | 5 | 50 | 2 | 0 | 0 | 3 |
|  | **Total** | **106** | **62** | **54** | **14** | **18** | **15** | **15** |
| LX283 (A1+B1 B1Δ) |  |  |  |  |  |  |  |  |
| □ |  |  |  |  |  |  |  |  |
| CBS6273 (A2 B2) |  |  |  |  |  |  |  |  |
|  | 1 | 12 | 3 | 25 | 0 | 1 | 1 | 1 |
|  | 2 | 14 | 0 | 0 | 0 | 0 | 0 | 0 |
|  | 3 | 13 | 7 | 54 | 0 | 0 | 1 | 1 |
|  | 4 | 11 | 7 | 64 | 5 | 2 | 0 | 0 |
|  | 5 | 11 | 3 | 27 | 2 | 1 | 0 | 0 |
|  | 6 | 12 | 3 | 25 | 3 | 0 | 0 | 0 |
|  | 7 | 12 | 3 | 25 | 0 | 0 | 0 | 0 |
|  | 8 | 12 | 7 | 58 | 3 | 0 | 4 | 4 |
|  | 9 | 14 | 9 | 64 | 6 | 0 | 3 | 3 |
|  | 10 | 14 | 7 | 50 | 6 | 0 | 1 | 1 |
|  | **Total** | **125** | **49** | **39** | **25** | **4** | **10** | **10** |
|  |  |  |  |  | A2+B2 B2Δ | A2+B2 B1 | A1 B2Δ | A1 B1 |
| CBS6039 (A1 B1) |  |  |  |  |  |  |  |  |
| □ |  |  |  |  |  |  |  |  |
| LX167 (A2+B2 B2Δ) |  |  |  |  |  |  |  |  |
|  | 1 | 8 | 2 | 25 | 1 | 0 | 0 | 1 |
|  | 2 | 8 | 3 | 38 | 1 | 1 | 0 | 1 |
|  | 3*** | 9 | 8 | 89 | 0 | 1 | 4 | 3 |
|  | 4*** | 9 | 7 | 78 | 3 | 0 | 1 | 3 |
|  | 5 | 12 | 9 | 75 | 3 | 0 | 0 | 6 |
|  | 6 | 8 | 5 | 63 | 3 | 1 | 0 | 1 |
|  | 7 | 12 | 12 | 100 | 0 | 3 | 9 | 0 |
|  | **Total** | **66** | **46** | **67** | **11** | **6** | **14** | **15** |
| CBS6039 (A1 B1) |  |  |  |  |  |  |  |  |
| □ |  |  |  |  |  |  |  |  |
| LX170 (A2+B2 B2Δ) |  |  |  |  |  |  |  |  |
|  | 1 | 11 | 8 | 73 | 2 | 0 | 3 | 3 |
|  | 2 | 14 | 8 | 57 | 5 | 1 | 2 | 0 |
|  | 3 | 13 | 10 | 77 | 3 | 1 | 6 | 0 |
|  | 4*** | 11 | 11 | 100 | 1 | 4 | 4 | 2 |
|  | 5*** | 14 | 13 | 93 | 4 | 0 | 0 | 9 |
|  | 6*** | 14 | 13 | 93 | 8 | 0 | 0 | 5 |
|  | 7 | 12 | 7 | 58 | 2 | 5 | 0 | 0 |
|  | 8 | 13 | 9 | 69 | 2 | 2 | 3 | 2 |
|  | 9 | 11 | 6 | 55 | 0 | 4 | 1 | 1 |
|  | 10 | 14 | 9 | 64 | 1 | 0 | 0 | 8 |
|  | 11 | 14 | 6 | 43 | 0 | 2 | 4 | 0 |
|  | 12*** | 14 | 10 | 71 | 2 | 2 | 1 | 5 |
|  | **Total** | **155** | **110** | **71** | **30** | **21** | **24** | **35** |

<sup>1</sup>: Basidia highlighted with asterisks (\*\*\*) are those from which basidiospores with representative genotypes were selected for whole genome sequencing and analysis.

We further assayed the mating abilities of a subset of progeny recovered from these tripolar crosses (Figures 2D and S5). As expected, progeny that inherited no *HD* alleles, such as those with *MAT* genotypes of “A2 B1Δ” or “A1 B2Δ”, were sterile when crossed with wild-type strains possessing compatible *MAT* alleles at the *P/R* (A) locus, because of the absence of heteroallelic HD-dimers in these crosses. In contrast, progeny harboring both a *MAT*-fused allele and a compatible *HD* allele from the *HD* locus (i.e. “A1+B1 B2” and “A2+B2 B1”) were self-fertile under mating inducing conditions, producing monokaryotic sexual hyphae with clamp cells, as well as basidia with abundant basidiospores (Figures 2D and S5), further supporting the conclusion that successful mating requires heteroallelic dimers formed by HD proteins of the opposite mating types. Additionally, the progeny of the “A1+B1 B2” and “A2+B2 B1” mating types are products of meiosis; thus, they have a recombinant nuclear genome (Figures S6), and consequently vary in their self-fertility, with some of them exhibiting more robust selfing than others (Figure S5), suggesting the presence of genetic elements outside of the *MAT* loci that contribute to the self-fertility of these progeny.

### Relocation of the *HD* locus affects mating in a bipolar *MAT* configuration

We observed significantly reduced sexual development in bilateral crosses of the *MAT*-fused strains, i.e. (A1+B1 B1Δ) × (A2+B2 B2Δ) crosses, with severely reduced production of sexual hyphae, basidia, and basidiospores (Figures 3A). This suggests that sexual reproduction is compromised in the bipolar matings between two *MAT*-fused strains. Because the *MAT*-fused strains showed normal fertility in unilateral tripolar crosses, we hypothesized that the bipolar mating defects were caused by compromised cell-cell fusion.

**Figure 3.**
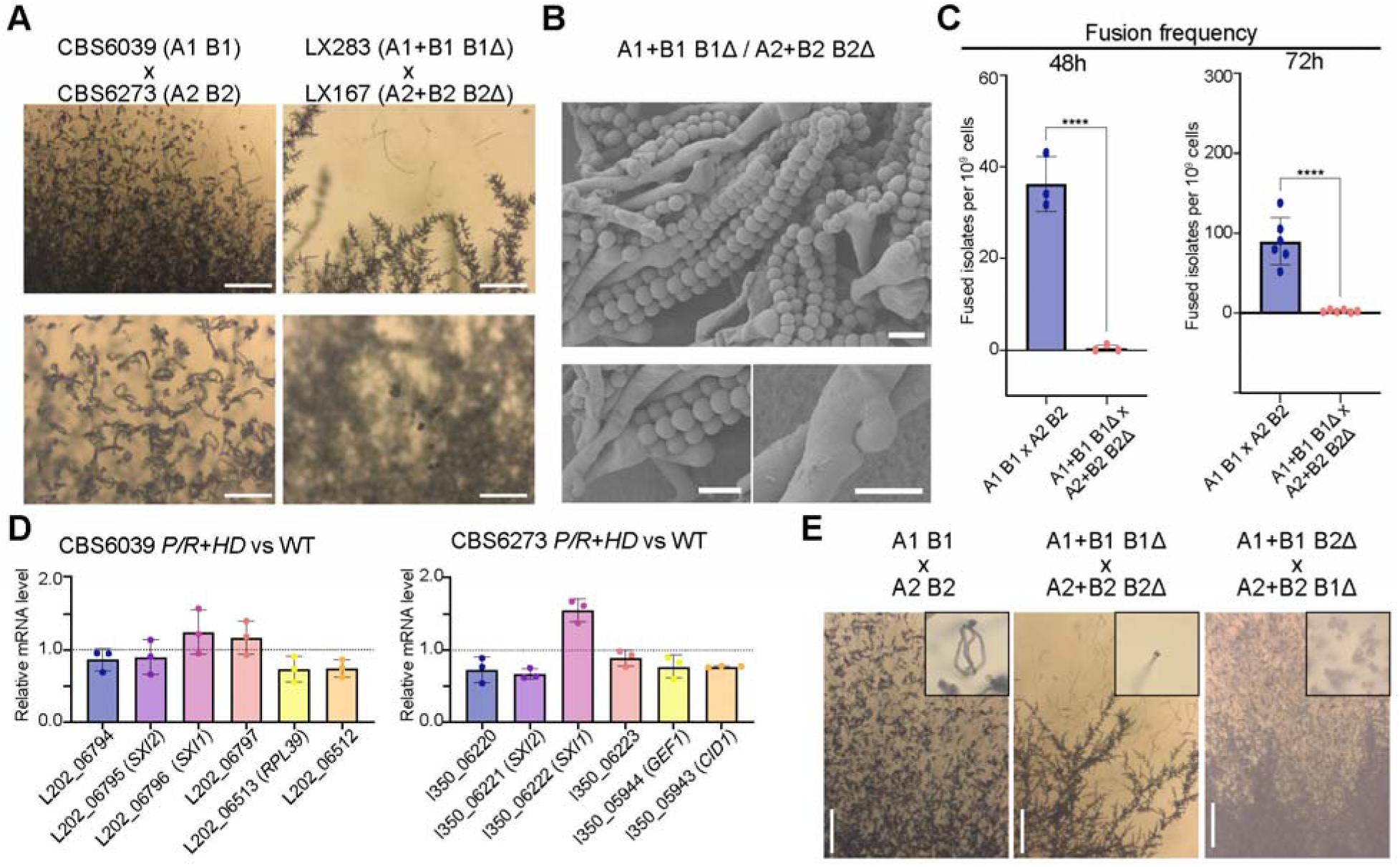
Relocation of the *HD* locus compromised mating in bipolar crosses. **(A)** Sexual hyphae production and sporulation observed in crosses between two *C. amylolentus* wild type tetrapolar isolates and their *P/R*+*HD* derived bipolar strains. Tested strains were cultured on V8 (pH=5) medium incubated in the dark at room temperature for 2 weeks. Upper panel, scale bar = 200 µm; lower panel: scale bar = 100 µm. (**B)** SEM revealed basidiospores attached to basidia and unfused clamp cells. Images were taken from the CBS6039 *P/R*+*HD* x CBS6273 *P/R*+*HD* selfing diploid isolate LX301, indicated as A1+B1 B1Δ / A2+B2 B2Δ. Scale bar: 5 µm (**C)** *HD* genes and their location are critical for fusion frequency. Each dot indicates one sample collected from one plate of 16 spots. Triplicates each were tested for 48 h and 6 samples each were tested for 72 h. Statistical analysis was performed with a one-way ANOVA, and asterisks indicate statistically significant differences. **** p-value < 0.0001. **(D)** Gene expression of *SXI1* and *SXI2* genes or their neighboring genes in *P/R*+*HD* vs WT. Graphs indicate relative mRNA level of target genes in the *P/R*+*HD* strain compared to wild type strain. RNA of three independent biological samples was extracted for qPCR determination. Relative gene expression was compared with the threshold cycle ΔΔCT method. Statistical analysis was performed with the T test. **(E)** Mating capability of F1 progeny isolated from selfing diploids. Tested strains were cultured on V8 (pH=5) medium incubated in the dark at room temperature for 2 to 3 weeks. Mating structures including sexual hyphae and basidiospores were assessed. Basidiospores were imaged at higher magnification (inset). Scale bar: 200 µm.

To test this, we estimated the frequencies of cell-cell fusion in bilateral crosses of *MAT*-fused strains and compared them with those in crosses between wild-type strains CBS6039 (A1 B1) and CBS6273 (A2 B2). The average fusion frequency between wild-type strains was ∼36 per 10 cells after 48 hours of co-culturing on V8 (pH=5) medium under mating inducing conditions, which increased to ∼90 per 10 cells after 72 hours of incubation. In contrast, the fusion frequencies in crosses of *MAT*-fused strains were only ∼0.42 per 10 cells after 48 hours and 2.56 per 10 cells after 72 hours, corresponding to an ∼86-fold and ∼35-fold reduction at the two time points, respectively (Figure 3C).

We next constructed diploid strains (LX301 and LX302) by fusing two *MAT*-fused strains of opposite mating types, LX283 (A1+B1 B1Δ) and LX167 (A2+B2 B2Δ), and tested their self-fertility under mating inducing conditions. These diploid strains with a homozygous *MAT*-fused configuration are self-fertile, undergoing robust sexual development and producing sexual hyphae and abundant basidiospores (Figure 3B), which are comparable to those produced by the dikaryotic strains formed during matings between the two wild-type isolates (Figure 1B).

Taken together, the reason that sexual development in the bilateral crosses of *MAT*-fused strains is highly reduced is likely due to a significant reduction in cell-cell fusion efficiency, rather than defects in subsequent stages of sexual development.

To determine if the defects in cell-cell fusion were due to altered gene expression, we conducted quantitative real-time PCR (qRT-PCR) analyses of the translocated *HD* genes as well as their neighboring genes in the *SH* loci in solo-cultures of the *P/R*+*HD* strains under mating inducing conditions compared to the wild-type strains. All of the tested genes had similar expression levels between the *MAT-*fused and the wild-type strains, suggesting that the deletion of the *HD* genes from their endogenous locus and their subsequent relocation into the *SH* loci did not significantly affect the expression of either the *HD* genes or their neighboring genes. These results suggested that decreased cell-cell fusion frequencies observed in bipolar crosses are unlikely due to transcriptional misregulation of the tested genes (Figure 3D). Instead, we hypothesize that the genetic relocation of *HD* genes may disrupt cis-regulatory or chromatin-dependent interactions that normally occur at the native *HD* locus, indirectly impairing early mating processes.

We next dissected basidiospores from 28 individual basidia (14 basidia each) from the two diploid strains with compatible *MAT*-fused alleles (LX301 and LX302). Of the 338 progeny dissected, 284 (84%) germinated, and the germination rates of the vast majority of basidia ranged between 50% and 100% (Table 2). The germination rates are higher than those observed in the tripolar crosses (Table 1). Among the 284 germinated progeny, two mating types (A1+B1 and A2+B2) were identified, suggesting the fused *P/R* and *HD* loci are segregating in their entirety during sexual reproduction. We further tested the fertility of 16 progeny recovered from these two selfing strains (Table 2). Similar to the progeny dissected from the tripolar crosses, we observed variation in the mating ability among progeny with identical *MAT* genotype, providing further evidence that genetic factors outside of the *MAT* loci are contributing to the fertility of *C. amylolentus* (Figure 3E).

**Table 2.** Genotyping analysis of spores dissected from bipolar crosses.

| Selfing diploid <sup>1</sup> | Basidia <sup>2</sup> | Dissected spores | Germinated spores | Germination rate (%) | Progeny mating-type |  |  |  |
| --- | --- | --- | --- | --- | --- | --- | --- | --- |
|  |  |  |  |  | A1+B1 B1Δ | A1+B1 B2Δ | A2+B2 B1Δ | A2+B2 B2Δ |
| LX301 | 1 | 14 | 8 | 57.14 | 3 | 0 | 0 | 2 |
|  | 2 | 14 | 13 | 92.86 | 6 | 1 | 5 | 1 |
|  | 3 | 11 | 9 | 81.82 | 1 | 5 | 2 | 1 |
|  | 4 | 12 | 10 | 83.33 | 0 | 2 | 4 | 4 |
|  | 5 | 12 | 11 | 91.67 | 0 | 4 | 4 | 3 |
|  | 6 | 14 | 13 | 92.86 | 5 | 3 | 4 | 1 |
|  | 7 | 12 | 12 | 100.00 | 3 | 0 | 0 | 9 |
|  | 8 | 12 | 10 | 83.33 | 8 | 0 | 0 | 2 |
|  | 9 | 11 | 11 | 100.00 | 1 | 0 | 0 | 10 |
|  | 10 | 13 | 13 | 100.00 | 8 | 0 | 0 | 5 |
|  | 11 | 11 | 4 | 36.36 | 3 | 0 | 1 | 0 |
|  | 12*** | 14 | 14 | 100.00 | 3 | 4 | 1 | 5 |
|  | 13*** | 11 | 11 | 100.00 | 3 | 4 | 1 | 3 |
|  | 14 | 12 | 7 | 58.33 | 4 | 0 | 2 | 1 |
|  | **Total** | **173** | **146** | **84.12** | **48** | **23** | **24** | **47** |
| LX302 | 1 | 11 | 11 | 100.00 | 6 | 0 | 0 | 4 |
|  | 2 | 11 | 8 | 72.73 | 3 | 0 | 1 | 4 |
|  | 3*** | 9 | 9 | 100.00 | 1 | 1 | 6 | 1 |
|  | 4 | 10 | 9 | 90.00 | 2 | 2 | 2 | 3 |
|  | 5 | 10 | 9 | 90.00 | 4 | 0 | 0 | 5 |
|  | 6 | 14 | 13 | 92.86 | 1 | 6 | 5 | 1 |
|  | 7 | 12 | 2 | 16.67 | 2 | 0 | 0 | 0 |
|  | 8 | 14 | 6 | 42.86 | 0 | 0 | 0 | 6 |
|  | 9 | 14 | 14 | 100.00 | 0 | 9 | 5 | 0 |
|  | 10 | 13 | 11 | 84.62 | 5 | 1 | 3 | 2 |
|  |  |  |  |  | A1+B1 B1Δ | A1+B1 B2Δ | A2+B2 B1Δ | A2+B2 B2Δ |
|  | 11 | 11 | 10 | 90.91 | 6 | 0 | 0 | 4 |
|  | 12*** | 11 | 11 | 100.00 | 4 | 2 | 2 | 3 |
|  | 13 | 13 | 13 | 100.00 | 1 | 0 | 0 | 12 |
|  | 14 | 12 | 12 | 100.00 | 0 | 6 | 6 | 0 |
|  | **Total** | **165** | **138** | **84.33** | **35** | **27** | **30** | **45** |

<sup>1</sup>: Both LX301 and LX302 are diploid strains generated by fusion of strains LX283 (CBS6039 *hdΔ::NAT shΔ::HD-HYG*) and LX167 (CBS6273 *hdΔ::NEO shΔ::HD-NAT*)
<sup>2</sup>: Basidia highlighted with asterisks (\*\*\*) are those from which basidiospores with representative genotypes were selected for whole genome sequencing and analysis. Sixteen progeny selected from these 4 basidia were also used for fertility tests.

### Progeny from tripolar and bipolar mating crosses underwent canonical meiosis

We have shown that among the progeny generated from “tripolar” and “bipolar” crosses, alleles at the *P/R* and *HD* loci underwent segregation, consistent with meiotic recombination. To further explore the genome-wide profiles of meiotic recombination, we conducted Illumina whole genome sequencing and variant analyses for 42 progeny recovered from these crosses involving engineered *MAT*-fused strains (Tables 1 and 2), which include: 1) 8 progeny from “tripolar” crosses between LX143 (A1+B1 B1Δ) and CBS6273 (A2 B2), comprising four representative progeny each from two tetratype (TT) basidia for the *P/R* and *HD MAT* loci (Table 1); 2) 18 progeny from the reciprocal “tripolar” crosses between CBS6039 (A1 B1) and LX167 or LX170 (both A2+B2 B2Δ), of which 14 were from four *MAT* TT basidia and 4 were from two *MAT* parental ditype (PD) basidia (Table 1); and 3) 16 progeny from selfing of the diploid strains LX301 and LX302, both of which are fusion products of *MAT*-fused haploid strains with compatible mating types (i.e. equivalent of “bipolar” crosses), with four representative progeny each from four independent basidia (Table 2).

The sequencing data were mapped to a *de novo* and improved nanopore-sequencing based telomere-to-telomere CBS6039 genome assembly in which 27 out of 28 telomere repeat ends are represented. Variant analyses provided clear evidence that canonical meiotic recombination occurred during these tripolar and bipolar crosses involving *MAT*-fused strains (Figure 4 and S6). Specifically, all of the progeny analyzed were recombinants of the two parental alleles, with crossovers observed throughout the genome (Figure S6). Additionally, analyses of four representative progeny from each basidia showed clear 2 : 2 balanced-inheritance of the two parental alleles across the genome, consistent with meiotic recombination and independent assortment. Furthermore, analyses of chromosome 10 (Figure 4) suggested there were on average 2 to 3 crossovers per chromosome, as well as suppressed crossovers in regions flanking the *P/R* locus, consistent with previous studies of meiotic recombination of *C. amylolentus* wild-type strains (28).

**Figure 4.**
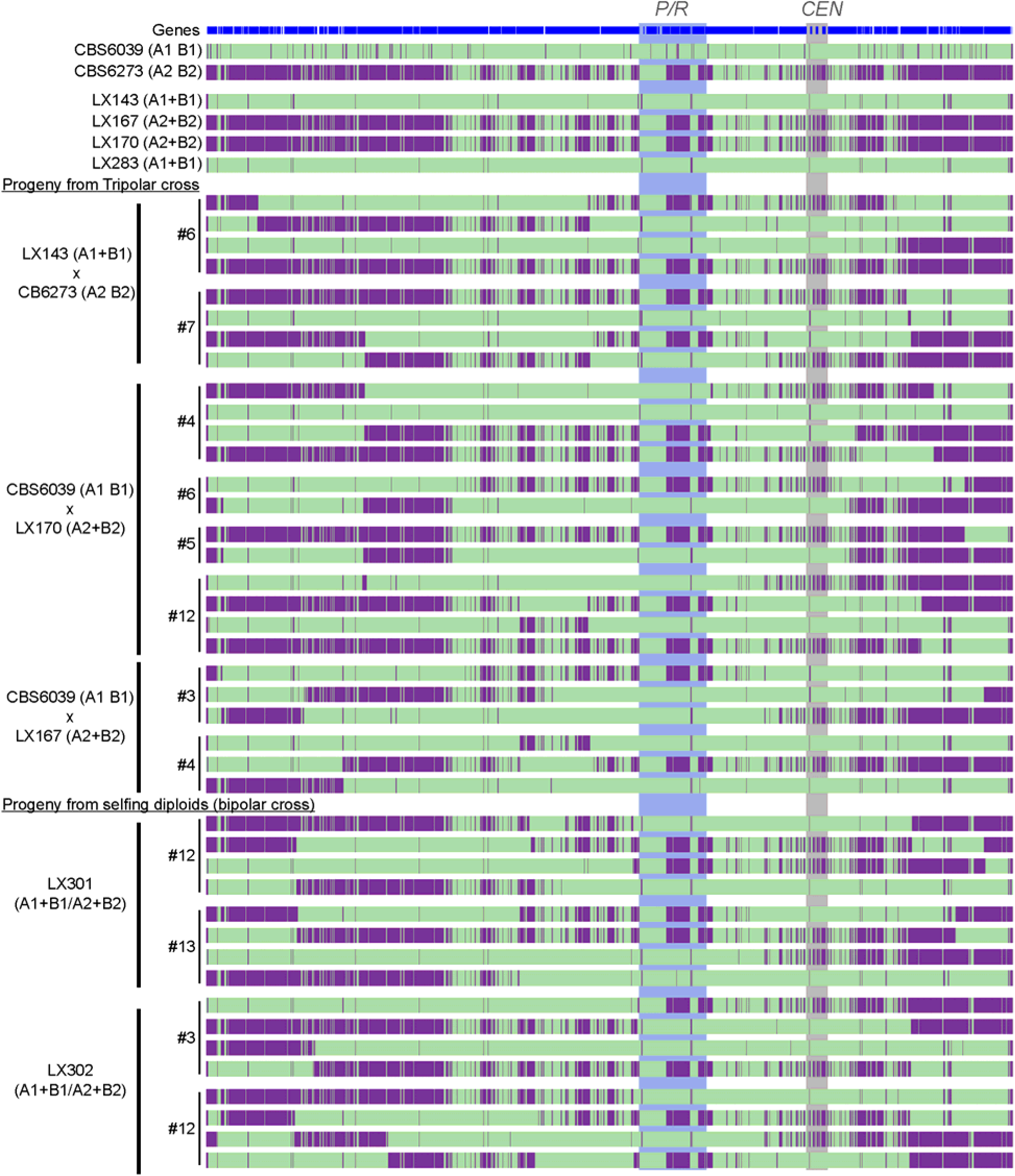
Variant mapping illustrates meiotic recombination in progeny from tripolar and bipolar crosses in *C. amylolentus*. Shown here are the results of SNP analysis of chromosome 10, on which the *P/R* locus is located. Illumina WGS reads of parental strains (wild type and engineered, top), as well as the progeny dissected from tripolar (middle) and bipolar (i.e. diploid selfing, bottom) crosses were mapped against the improved CBS6039 reference genome assembled using the Nanopore long-read sequencing data. Green indicates sequences identical to CBS6039, while purple indicates variants from the CBS6273 genome. The progeny from the same basidium were grouped together, with their basidium number, and the cross from which they were dissected listed on the far left and correspond to those listed in Tables 1 and 2. The *P/R* locus and centromeric (*CEN*) regions are highlighted in shades.

Taken together, our studies suggest that the relocation of the *HD* genes does not affect meiosis, and canonical meiotic recombination occurred during these tripolar and bipolar crosses in *C. amylolentus*.

### Sxi1/Sxi2 induces DNA replication genes during sexual development

To determine the gene expression impact of the HD transcription factors (TFs), we isolated fused cells from the mating crosses between the two wild-type isolates and the two *hd*Δ mutants, respectively, and then performed RNA Seq on two isolates— one wild-type WT/WT (LX44) and one *hd*Δ/*hd*Δ mutant (LX230). These two isolates were analyzed instead of mating cocultures to minimize background, as *C. amylolentus* shows self-filamentation, a low frequency of cell-cell fusion, variable mating phenotypes, and *hd*Δ mutants fail to form sexual structures. This design enabled direct comparison of the Sxi1-Sxi2 response in WT/WT, which develops clamp cells and sexual hyphae, versus mutant, which remains vegetative. RNA samples were extracted from *C. amylolentus* diploid cells grown on V8 (pH=5) agar medium and YPD agar medium for 48 hours (Figure 5A).

**Figure 5.**
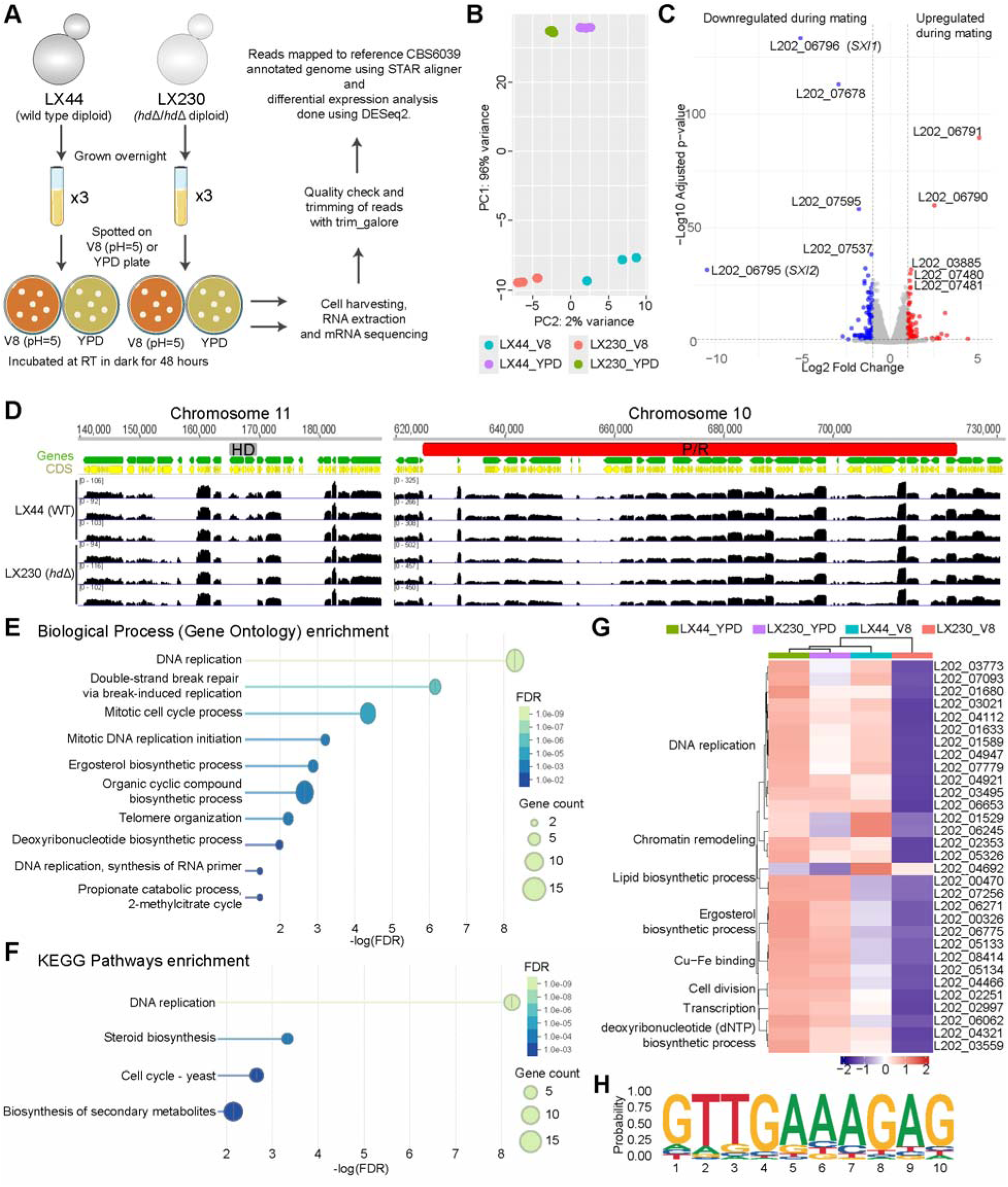
Impact of HD protein heterodimers during mating. **(A)** Schematic workflow of RNA-seq experiment, including sample collection, RNA extraction, library preparation, sequencing, and downstream bioinformatic analysis. **(B)** Principal component analysis (PCA) plot showing sample clustering based on the first two principal components (PC1 and PC2). Three biological replicates of the wild-type and the mutant diploids under YPD and V8 (pH=5) media conditions were analyzed by RNA sequencing. **(C)** Volcano plot highlighting differentially expressed genes (DEGs) between mutant and WT, with significantly up (red) and down-regulated (blue) genes indicated. **(D)** Mapping of RNA-seq reads to the reference genome (CBS6039). Read coverage tracks display the distribution and abundance of aligned reads across genomic regions of the *P/R* and *HD* locus. Differences in read density reflect relative transcript expression levels. No reads corresponding to *SXI1* or *SXI2* were detected in the *hd*Δ diploid, confirming successful deletion. The neighboring genes of *SXI1* and *SXI2*, as well as other genes within *P/R* loci displayed similar expression patterns between the mutant and WT. **(E)** GO term enrichment analysis identified significantly enriched biological processes associated with DNA replication and ergosterol biosynthesis among down-regulated differentially expressed genes (DEGs). **(F)** KEGG pathways enrichment analysis of the down-regulated DEGs of mutant vs WT. **(G)** Heatmap representation of expression profiles for the 31 DEGs across samples on both YPD and V8 agar media. Mean of triplicates is presented here. Categories of GO enrichment are denoted by left. **(H)** One predicted biding motif that is specifically enriched in the down-regulated genes and present in the promoter regions of all genes listed in (G).

RNA-seq reads were mapped to the reference CBS6039 genome and gene expression levels were measured as gene counts followed by DESeq2 analysis to identify significantly differentially regulated genes (log_2_ fold change >1 and padj<0.05). PCA of transcriptomic profiles showed separation between the biological replicates from YPD and V8 media conditions, highlighting that transcriptomic profile changes dramatically between these two media conditions. PC2 reflected the variation between the WT/WT (LX44) and *hd*Δ/*hd*Δ (LX230) diploids with more separation between the sample groups of the V8 condition as compared to the YPD media, suggesting a more prominent impact of *hd* deletion in mating condition (Figure 5B).

A pair-wise comparison between the wild type and mutant strains revealed that 18 genes were differentially expressed (10 up- and 8 down-regulated) in the mutant as compared to the wild type on YPD agar medium, whereas 168 genes were differentially regulated (80 up- and 88 down-regulated) under mating conditions (Figure 5C and Table S2). Among these, 7 genes were shared between YPD and V8 media conditions. As expected, the *SXI1* (L202_06796) and *SXI2* (L202_06795) were the two most down-regulated transcripts in both conditions (Figure 5C, 5D, and Table S2). Interestingly, two genes at the boundary of *HD* locus (L202_06790 and L202_06791) were up-regulated in the mutant under both conditions, even though the expression of immediate *HD*-neighboring genes remained unchanged, consistent with qRT-PCR results (Figure 3D, and Table S2). These results suggest that *HD* deletion activates specific upstream genes, whose functional relevance remains undetermined, without broadly affecting neighboring genes, implying a potential cis-regulatory or chromatin boundary function at the *HD* region. Among the remaining three shared genes, two (L202_07283, L202_07678) code hypothetical proteins, and one (L202_07595) encodes a condensation domain-containing protein; however, their roles in sexual reproduction remain to be elucidated.

GO enrichment as well as STRING network analysis failed to identify any group of significance among the differentially regulated genes under the YPD condition as well as among the up-regulated genes under the V8 condition. Interestingly, analysis of down-regulated genes on V8 medium during mating revealed significant enrichment of genes involved in DNA replication—including all six MCM complex subunits (*MCM2-7*) and dNTP biosynthesis genes, and genes involved in lipid biosynthesis, specifically ergosterol biosynthesis (Figure 5E—G). Further analysis of expression profiles of these genes across all conditions revealed that *hd* deletion caused down-regulation of some of the DNA replication genes even under the YPD condition (fold change 1.3-1.4) (Figure 5G and Table S2). Furthermore, genes involved in lipid biosynthesis are similarly down-regulated in the wild-type when grown on V8 agar medium, but this down-regulation is exacerbated in the mutant. Both analyses suggest a more sexual reproduction specific role for Sxi1 and Sxi2.

Motif analysis of promoter regions among these genes identified several motifs in each analyzed set (up-regulated, down-regulated, or combined) (Figure S7). Interestingly, one of these motifs was specifically enriched in the down-regulated gene sets and was present in the promoter of all genes involved in DNA replication and lipid biosynthesis (Figure 5H), suggesting a cis-regulation of these genes through Sxi1-Sxi2 binding. We hypothesize that mutants lacking Sxi1 and Sxi2 fail to replicate their genome properly, which results in failure to undergo post-fusion stage of sexual cycle. Importantly, these mutants do not exhibit a defect during vegetative growth, suggesting the impact on replication is sexual reproduction specific. Furthermore, an impact on lipid biosynthesis could further exacerbate the defect during sexual cycle. Overall, these results show that HD transcription factors orchestrate a multi-layered program integrating DNA replication, cell-cycle progression, and lipid and sterol biosynthesis to establish and sustain hyphal growth during sexual reproduction.

## Discussion

The tetrapolar mating system, defined by two unlinked *MAT* loci (*P/R* and *HD*), is a distinctive feature of basidiomycetes, in contrast to the bipolar *MAT* configurations of ascomycetes and zygomycetes. In this study, by mutational analysis of the *HD* genes, we affirmed that the *MAT* homeodomain (HD) proteins function as key regulators of sexual development in *C. amylolentus*, with at least one non-self Sxi1–Sxi2 heterodimer required and sufficient to initiate dikaryotic growth and reproductive progression. Transcriptome analyses revealed that these heterodimers activate DNA replication and lipid biosynthetic programs during mating, underscoring their potential roles in coupling metabolic and developmental transitions. By relocating the HD transcription factors into the *P/R MAT* locus, we constructed *C. amylolentus* strains with fused *MAT* loci, and successfully established three modes of sexual reproduction: the ancestral tetrapolar, the transitional tripolar, and the derived bipolar (Figure 6). These results demonstrate that the tetrapolar system of *C. amylolentus* is capable of transitioning to a bipolar configuration, providing evidence that mating systems can evolve through stepwise genomic reorganization of *MAT* loci. More broadly, our findings highlight the plasticity of fungal sexual systems and suggest that the chromosomal position of the *HD* genes is a critical determinant of sexual identity, with wider implications for understanding how genomic architecture shapes the emergence and diversification of reproductive strategies across fungi.

**Figure 6.**
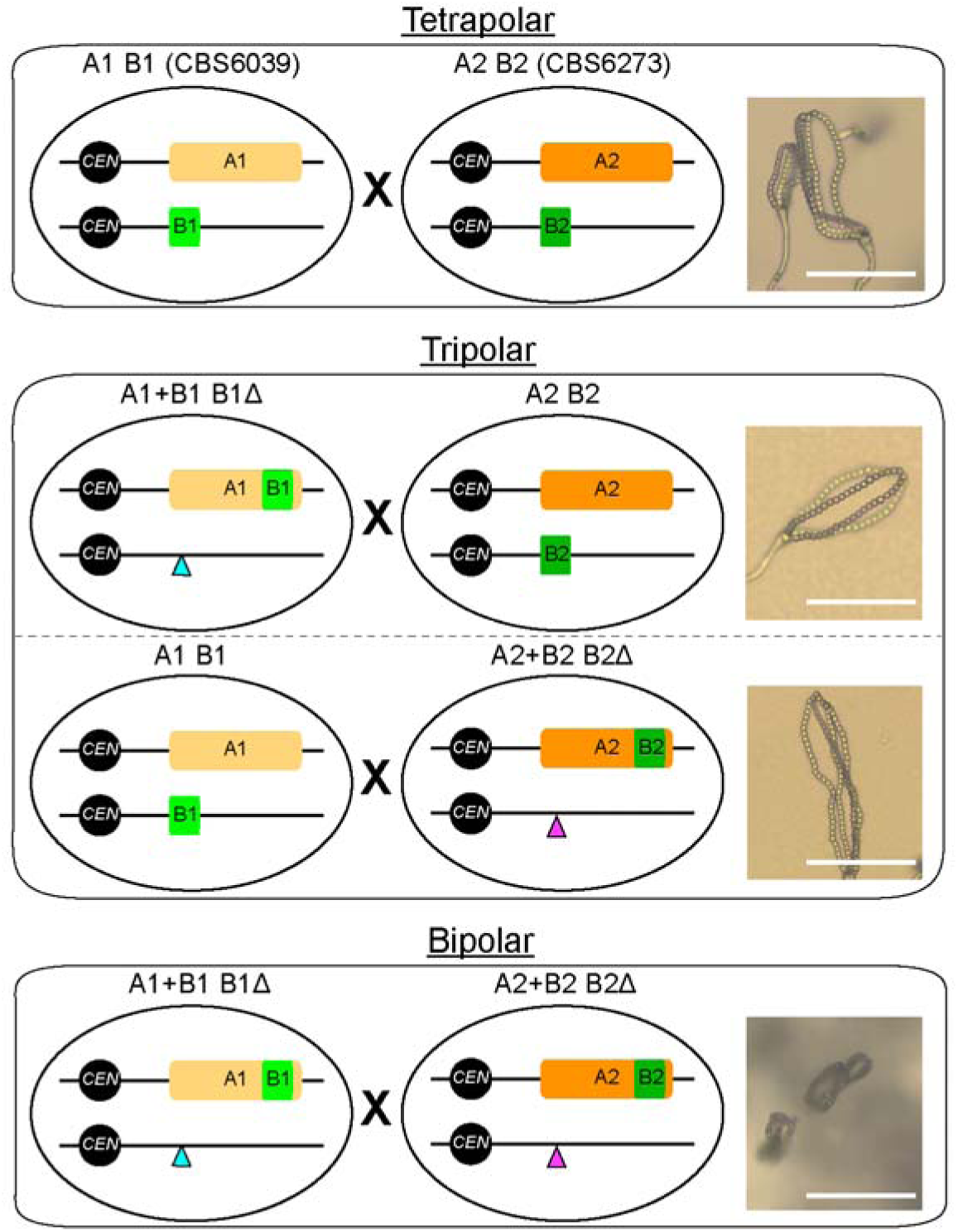
Diagrams depicting *MAT* loci configurations in the tetrapolar, tripolar, and bipolar crosses. In the two *C. am*ylolentus natural isolates, the two *MAT* loci, *P/R* and *HD*, are located on chromosomes 10 and 11, respectively. Thus, mating between CBS6039 (A1 B1) and CBS6273 (A2 B2) is tetrapolar (top). With the CRISPR-Cas9 system, we deleted the *HD* genes from their endogenous location (the blue and purple triangles) and then relocated them into the *P/R* locus (the light and dark green rectangles) in both CBS6039 and CBS6273 backgrounds, generating two *P/R*-*HD* linked strains, “A1+B1 B1Δ” and “A2+B2 B2Δ”. Crosses between these *MAT*-engineered strains are equivalent of bipolar mating (bottom), and crosses between each of these *MAT*-engineered strains and the natural isolate with compatible mating type are then considered to be intermediate tripolar (middle). Images of basidia and basidiospore chains produced from each type of mating are shown in the right (scale bar: 50 µm).

### HD proteins – conserved regulators of fungal reproduction

HD proteins act as conserved regulators of fungal reproduction, coupling partner recognition with developmental progression (6, 33, 44). In *C. amylolentus*, the non-self heterodimers of the HD proteins are required to initiate mating and maintain the dikaryotic state, with a single pair of Sxi1-Sxi2 sufficient to trigger sexual development (Figure 1A and 1B). Our modeling with AlphaFold suggests Sxi1 and Sxi2 of the same strain might also form a heterodimer, although no selfing has been observed in *C. amylolentus*. It is possible that only one of the two HD genes in each strain is expressed under mating inducing conditions, preventing the formation of self-HD-dimers. Alternatively, AlphaFold may not have sufficient discriminatory power to detect authentic *in vivo* specificity.

We found that HD-mutant diploids failed to undergo sexual development under mating inducing conditions. Transcriptome profiling revealed that the *MCM2-7* complex genes, central to DNA replication licensing, were markedly down-regulated in mutants, indicating replication and cell-cycle defects may occur in the absence of the Sxi1–Sxi2 complex. In eukaryotes, the Mcm2–7 proteins are universally essential for replication initiation, S-phase entry, and broader nuclear processes including chromosome segregation, histone inheritance, DNA damage sensing, and transcription (45, 46). Thus, our RNA-seq analysis may uncover a previously unrecognized role for Sxi1-Sxi2 complex in promoting DNA replication. Further analysis of the promoters identified a down-regulated gene-specific motif (Figure 5H) that was present in all replication-related genes. This analysis also identified another motif that is present in more than 60% of down-regulated genes and resembles the conserved MBF consensus binding motif (Figure S7). MBF (Mbp1/Swi6) is a well-known sequence-specific transcription factor that activates gene expression during the G1/S transition of the cell cycle in yeast (47, 48). These findings suggest that the Sxi1–Sxi2 complex may cooperate with the MBF-like transcription factor in *C. amylolentus* to coordinate the activation of DNA replication and other G1/S-associated genes during sexual development. Future work should determine whether the uncharacterized upstream genes interact with the *HD* locus and identify direct HD binding targets via ChIP-seq or motif-binding assays to clarify the underlying regulatory network.

These Sxi-induced replication-related genes do not contain canonical HD-binding motifs, suggesting indirect regulation via noncanonical elements or cofactors. Supporting this, in *U. maydis*, bE/bW-dependent induction of *IGA2* requires additional regulatory sequences (49), and bE(ts) strains display multinucleate hyphal tips, suggesting failed nuclear division (50). Additionally, in *Ustilago maydis*, the bE/bW heterodimer induces G2 arrest and filamentous growth (13); while in *Coprinopsis cinerea*, mating triggers genes involved in cell-cycle regulation, cytoskeleton dynamics, and cell wall biogenesis, with clamp cell formation blocked by DNA replication inhibitors (51–53). Our RNA-seq analysis suggests that the Sxi1–Sxi2 complex may regulate lipid biosynthesis processes, and fungal ergosterol is critical for sexual reproduction and sporogenesis in *C. neoformans* (54). Taken together, our observations suggest that Sxi1–Sxi2 may couple DNA replication with morphogenesis to coordinate sexual development and reveal expanded functions of the HD1–HD2 heterodimer that encompass DNA replication, metabolism, and sexual differentiation, possibly to ensure synchronized nuclear division, morphogenesis, and cellular homeostasis during development.

### Mating-system transitions and evolution

Transitions from tetrapolar to bipolar mating systems have occurred repeatedly across basidiomycetes, often through independent evolutionary routes (55, 56). In smut fungi, Bakkeren and Kronstad (57) initially showed that the bipolar mating-type locus of *Ustilago hordei* arose through fusion of the *a* and *b* loci into a single nonrecombining region with two alleles, mediated by chromosomal translocation (58). Evolutionary mechanisms in other basidiomycetes remain largely unexplored. In *Cryptococcus*, ancestral tetrapolar species such as *C. amylolentus* maintains unlinked, recombining *MAT* loci, and rearrangements around the *P/R* locus indicate the potential for structural evolution (28, 59).

Our functional study reveals several key insights into how mating-type (*MAT*) locus organization influences sexual development in *C. amylolentus*. First, engineered bipolar mating efficiency was reduced relative to its ancestral tetrapolar system, highlighting the critical influence of *HD* locus positioning on sexual competency. This could also serve as a functional “barrier” for the complete transition between tetrapolar and bipolar, maintaining the integrity of each mating system. Second, deletion of *HD* genes altered expression of nearby upstream genes without affecting immediately adjacent loci, suggesting that the *HD* locus may act as a regulatory boundary, maintaining local chromatin organization and insulating local gene expression. While most boundary genes near the *P/R* and *HD* loci are dispensable for mating, exceptions such as *STE20* are essential for maintaining sexual hyphae polarity (60), indicating that subtle regulatory shifts can indirectly influence mating efficiency. Third, while bipolar mating crosses exhibited delayed mating, the recombinant progeny derived from these crosses displayed enhanced fertility. This enhancement may simply reflect the recombinant nature of these meiotic progeny, in which favorable allele combinations outside of *MAT* improve fertility. Alternatively, these findings suggest the existence of genetic elements outside the *MAT* loci that can enhance mating and facilitate the transition to bipolar mating system.

We observed pronounced cell-cell fusion defects between A1+B1 and A2+B2 strains, as well as between two *hd*Δ mutants. We ruled out the possibility that this defective phenotype was due to artifacts introduced during genetic manipulation by whole genome sequencing (Figure S6) as well as by analyzing multiple independent strains. We hypothesize that the impaired fusion mainly arises because the functional roles of HD were not fully restored after the relocation of *HD* genes to the *P/R* locus. A possible explanation is that *HD* deletion disrupted local regulatory elements required for early mating responses. RNA-seq profiling supports this view, revealing *HD* deletion resulted in up-regulation of neighboring genes of unknown function and down-regulation of genes involved in DNA replication and lipid biosynthesis. These transcriptional changes likely compromise overall cellular activity and membrane dynamics necessary for efficient fusion. Given that DNA synthesis supports cell cycle progression and lipid remodeling mediates membrane merger, incomplete restoration of Sxi1 and Sxi2 function after relocation likely underlies the persistent fusion defect. Recent findings in *C. deneoformans* provide a relevant parallel: Sxi1α has been shown to inhibit cell fusion during unisexual reproduction, whereas its partner, Sxi2**a**, exerts only limited inhibition under specific genetic contexts (61). These observations raise the possibility that homeodomain proteins may modulate fusion competence in a context-dependent manner. Taken together, our findings expand the role of HD1–HD2 heterodimers beyond dikaryotic hyphal growth, showing that HD proteins integrate mating identity, metabolism, and fusion competence, and thereby link molecular function to evolutionary constraints on *MAT* locus architecture and mating system transitions.

Fungal mating systems display remarkable diversity, even among closely related species. Here, we provide the first mechanistic evidence that fusion of the *MAT* loci can drive the evolutionary transition from a tetrapolar to a bipolar mating system, with the genomic position of the *HD* locus playing a pivotal role in this process. Our findings reveal both the inherent flexibility enabling transitions between distinct mating configurations and the potential presence of intrinsic constraints that preserve the integrity and stability of each system.

## Materials and methods

### Strains, media and culture conditions

A full list of strains analyzed in this study is in Table S1. Strain stocks were maintained in 15% glycerol at −80°C. Prior to experiments, strains were grown on YPD agar (1% yeast extract, 2% Bacto peptone, 2% dextrose, 2% Bacto agar) at room temperature (22°C) for 24 to 48 hours unless stated otherwise. Cells were grown in 5 ml of liquid YPD medium (1% yeast extract, 2% Bacto peptone, 2% dextrose) at room temperature with rotation at 75 rpm. Transformants were selected on YPD plate containing 200 μg/ml nourseothricin (clonNAT, Gold Biotechnology), 100 μg/ml neomycin/G418 (Sigma N1687) or 200 μg/ml Hygromycin B (Sigma SBR00039). Basidia-specific spore dissections were performed after 2 weeks of mating, and the spore germination rate was scored after 4 days of dissection. Powder DTT (Sigma D0632) and W-7 hydrochloride (Sigma 681629) was dissolved in DMSO (Sigma 472301) to make stocks of DTT (1 M) and W-7 hydrochloride (5 mg/ml).

### AlphaFold modeling

AlphaFold modeling was done using AlphaFold3 available online (https://alphafoldserver.com/). The models were generated in various combinations and configurations for CBS6039 Sxi1, CBS6039 Sxi2, CBS6273 Sxi1, and CBS6273 Sxi2. The protein sequences were used to predict each of the homodimer and heterodimer configurations. For each prediction, 5 different models were obtained from the AlphaFold server, and only model_0.cif was used for analysis and presentation. All generated models were processed using ChimeraX 1.9 for presentation purposes in Figure S1 (62, 63).

### Genomic DNA extraction

Genomic DNA of the reference wild-type strains *Cryptococcus amylolentus* CBS6039 and CBS6273 was extracted using a CTAB protocol as described previously (64). Genomic DNA examined in this study was extracted using a mini-preparation protocol (65). Briefly, cells were collected from overnight culture (about a 2 cm patch) on YPD plate and suspended in 300 μl of lysis buffer (100 mM Tris pH=8.0, 50 mM EDTA, and 1% SDS). Cells were incubated at 65°C for 45 min with vortexing very 15 minutes then chilled in ice for 5 min. Next, 150 μl of protein precipitation buffer was added to the cell suspension. Cells were then vortexed for 30 sec followed by centrifugation at 14,000 rpm for 10 min. The carefully removed 400 μl of supernatant was mixed with 400 μl of isopropanol (Sigma). DNA was precipitated by isopropanol at room temperature over 10 min. Finally, the precipitated DNA samples were washed with 500 μl of 70% ethanol, air dried, and dissolved in 100 μl of sterile water or TE buffer.

### The Cas9, the sgRNA, and the gene deletion cassettes construction

To generate a *C. amylolentus* Cas9 expression construct, we modified a plasmid pXL1 (66) by using native promoter and terminator of *C. amylolentus TEF1*. Briefly, the native promoter and terminator sequences of *TEF1* (L202_04966) were amplified from *C. amylolentus* CBS6039 strain (all primers used in this study are listed in Table S1). The PCR product was confirmed by agarose gel and then purified using the Qiagen Gel Extraction and Purification Kit (Cat no. 28115, Qiagen, Germany). *TEF1* promoter sequence was digested with XbaI and FseI restriction enzymes, and *TEF1* terminator sequence was digested with ApaI and PacI restriction enzymes. The digested product was ligated into the plasmid pXL1 by replacing the previous *GPD1* promoter and the *GPD1* terminator with the NEB Quick-ligation kit (NEB M2200S). The resulting plasmid pLX1-Cas9 served as the template to amplify the whole *C. amylolentus* Cas9 expression cassette for transformation (Figure S2).

The sgRNA construct was generated as described previously (66). Briefly, the native U6 promoter was amplified from a *C. deneoformans* strain JEC21 while the scaffold of sgRNA with 6-T terminator was amplified from the plasmid pYF515 (66). The U6 promoter, 20-nt guide sequence, scaffold of sgRNA were assembled in the order shown in Figure S2A by a single-joint PCR (67). To optimize the expression of drug selection markers, we adapted native promoter and terminator of *C. amylolentus ACT1* (L202_07304) to the nourseothricin acetyltransferase gene *NAT1* expression cassette (*NAT*) and neomycin resistance gene expression cassette (*NEO*). Briefly, the native promoter and terminator sequences of *ACT1* were amplified from genomic DNA of *C. amylolentus* CBS6039. Then the drug selection sequences amplified from the plasmid pSDMA25 (*NAT*) and pSDMA57 (*NEO*) (68) then fused with promoter and terminator of *C. amylolentus ACT1* by overlapping PCRs. The fused fragment was then ligated to a vector pPCR2.1 using a TOPO™ TA Cloning system (Invitrogen, Cat no 450641). Electroporation to CBS6039 competent cells confirm that *C. amylolentus NAT* and *NEO* expression cassettes confer resistance to NAT and G418, respectively.

Homologous flanks of the targeted genes were amplified from the genome of the wild type strains: CBS6039 and CBS6273. DNA repair constructs were prepared in two different ways depending on their sizes. We constructed deletion cassettes of individual *SXI1* or *SXI2* or the combination of both. For mutant construction in CBS6039, the fragments of 5’ homology, selection marker, and 3’ homology were amplified, purified and then fused by a double-joint PCR (69, 70). For mutant construction in CBS6273, the three fragments were fused by using NEBuilder® HiFi DNA Assembly master mix, then amplified with the resulting plasmid DNA as template. DNA assembly product was transformed to NEB 10 beta competent *E. coli* cells (Cat no. C3019H). Schemes of deletion cassettes are shown in Figure S2B.

For each strain construction, three DNA products, including the Cas9 cassette, the sgRNA cassette, and the gene deletion or *HD* relocation cassette, were prepared as described above and eluted with ultrapure water. The DNA mixture of three DNA products was then concentrated by speed vacuum when necessary and introduced by electroporation as described below.

### Electroporation in Cryptococcus amylolentus

Electroporation was performed as we described previously with modifications (66, 70, 71). Briefly, freshly revived *C. amylolentus* cells were incubated in 5 ml of YPD liquid medium. Cells were grown overnight at room temperature with rotation at 75 rpm. The overnight culture was then transferred into 50 ml of fresh YPD medium with an initial inoculum of OD_600_ = 0.3. Cells were cultured for additional 4 to 5 hours until the cell density reached OD_600_ between 0.9 and 1.0. After treated in hydroxyurea (50 mM) for extra 1.5 to 2 hours, cells were pelleted by centrifugation at 3220 xg for 5 min at 4 °C, washed twice with ice-cold water, then suspended in 10 ml of ice-cold electroporation buffer (EB) buffer (10 mM Tris-HCl pH=7.5, 1 mM MgCl_2_, and 270 mM sucrose) with 1 mM DTT and 5 mg/ml W-7 hydrochloride. Finally, cells were harvested by centrifugation and were resuspended in EB buffer (150 μl) after 2 hours of incubation on ice. For each transformation reaction, 45 μl of cell suspension was mixed gently with 5 μl DNA mixture then added to a precooled 2 mm gap electroporation cuvette. Electroporation was done with a Bio-Rad MicroPulser Electroporator in the bacterial mode (V = 0.45 kV with t optimized for 5 msec). The electroporated cells were then suspended in 1 ml of YPD medium added 5 mg/ml W-7 and cultured at room temperature for 2.5 hours before being plated onto the appropriate selective agar medium (YPD with additional NAT or G418 or Hyg). Transformants were counted and patched to fresh selective media after 3 days of incubation at room temperature.

### Mutant construction

Deletion mutants of *HD* genes were made in wild type CBS6039 and CBS6273. Cas9 cassette was amplified from plasmid pLX-1 with primer set LX202 and LX203. sgRNAs were made to target *SXI1* and *SXI2* (named as *YFG1*) in this study. Deletion cassettes were amplified by PCR (Figure S2B). For each transformation, competent cells were transformed with approximately 1 μg of Cas9 DNA cassette, 0.5 μg of sgRNA DNA cassette, 2 μg of deletion cassette DNA. Transformants were selected on YPD+ NAT or YPD+ G418 agar plates. Candidate transformants were genotyped by PCR to validate deletion of the *YFG1* ORF and integration of the selection marker at the *YFG1* locus (Figure S2A). Primer sets used for PCR genotyping validation were indicated on the top of each gel image. Resultant mutant strain genotypes were *yfg1*Δ::*NAT* in CBS6039 and *yfg1*Δ::*NEO* in CBS6273. Following this strategy, mutants of *sxi1*Δ::*NAT, sxi2*Δ::*NAT,* and *sxi1*Δ *sxi2*Δ::*NAT* were generated in CBS6039, and mutants of *sxi1*Δ::*NEO, sxi2*Δ::*NEO,* and *sxi1*Δ *sxi2*Δ::*NEO* were generated in CBS6273 (Figure S2C).

To relocate the *HD* genes adjacent or within the pheromone/receptor *MAT* locus, we screened for one potential “safe haven” (SH) region within the boundary of *P/R* locus in CBS6039. To construct the *SH* deletion (*SH*-Marker) in the wild type CBS6039, the *SH* deletion cassette was generated by Gibson assembly and then amplified from the plasmid template by primers LX313 and LX314. To construct the *HD* locus relocation in the CBS6039 *hd*Δ strain background, the *HD* relocation cassette was assembled and then purified from the plasmid by using restriction enzymes EcoRI and XbaI and the QIAEX II System Gel Extraction Kit (Cat no. 20021, Qiagen, Germany). Then 173 bp of *SH* was replaced by a *NAT* marker in CBS6039 or by *HD* genes and a *NEO/HYG* marker in CBS6039 *hd*Δ with Cas9 DNA cassette and sgRNA DNA cassette targeting the *SH* region. Strain CBS6039 *hd*Δ was transformed with approximately 1 μg of Cas9 DNA cassette, 0.5 μg of sgRNA DNA cassette, 4 μg of *HD* relocation cassette. Transformants were selected on YPD+ NAT+ G418 or YPD+ NAT+ HYG agar plates. Candidate colonies were further genotyped by PCR to confirm the *HD* genes were inserted into indicated locale in the *P/R* locus. LX143, and LX283 were validated to be CBS6039 *P/R+HD* strains (Figure S3).

Similarly, a potential SH within *P/R* locus was chosen for *HD* locus relocation in CBS6273. The *SH* deletion cassette was assembled and then amplified from the plasmid template with primers LX315 and LX316. With our CRISPR-Cas9 system, 261 bp of the *SH* was replaced by a *NEO* marker in CBS6273 or by the *HD* genes and a *NAT* marker in CBS6273 *hd*Δ. The *HD* relocation cassette was assembled and then purified from the plasmid with restriction enzymes AleI-v2 and EcoRV. Transformants were selected on YPD+NAT+G418 agar medium. Candidate colonies were further genotyped by PCR. LX167 and LX170 were validated to be CBS6273 *P/R+HD* strains (Figure S3).

### Mating assays

Mating plates were set up as described previously (72). Sexual reproduction assays were performed on V8 (pH=5) plates for 2 to 3 weeks prior to imaging and spore dissection. Basidiospores generated during sexual reproduction were randomly dissected and plated on YPD plates for 2 to 3 days to determine spore viability and germination rates. Germinated progeny were further spotted on YPD plates with and without drug(s) to differentiate their mating types (segregation of chromosome 10 and 11) based on gene deletion/relocation alleles marked with dominant drug resistance markers. Specifically, NAT^R^ indicates either the deletion of the *HD* genes (B1Δ) on chromosome 11 of CBS6039 or the fused *P/R+HD* locus (A2+B2) on chromosome 10 of CBS6273; NEO^R^ indicates the deletion of the *HD* genes (B2Δ) on chromosome 11 of CBS6273, or the fused *P/R+HD* locus (A1+B1) on chromosome 10 of CBS6039; HYG^R^ indicates fused *P/R+HD* locus (A1+B1) on chromosome 10 of CBS6039; and non-resistance indicates the wild type A1 B1 or A2 B2 (Figure S4).

### Fusion frequency assays

In *C. amylolentus* sexual reproduction, LX21 (CBS6039 *NAT*) and LX26 (CBS6273 *NEO*) were used as genetically marked wild type strains to study the fusion competency of *P/R*+*HD* strains. Strains for each fusion pair were grown overnight on YPD agar plate at room temperature. Cells were collected and suspended in dH_2_O and diluted to a final density of OD_600_= 2. Then, 5 μl of equal-volume mixed cells were spotted on V8 (pH=5) medium (∼16 spots each plate) and incubated face up for 48 hours or 72 hours in the dark at room temperature. The cells from one plate were then harvested, resuspended with dH_2_O, and plated in serial dilution on both YPD medium and YPD medium supplemented with NAT, G418, and/or Hygromycin according to the selection markers in the tested strains. The cells were incubated for five days at room temperature. Cell-cell fusion frequency was measured by counting the average number of double drug resistant cfu/total cfu.

### Light and scanning electron microscopy (SEM)

Mating inducing media plates with mating structures (hyphae, basidia and basidiospores) were imaged with an AxioScop 2 fluorescence microscope. SEM samples were prepared following a previously described protocol (73), with minor modifications. Briefly, a ∼2 cm × 2 cm V8 agar plug containing mating structures was fixed in PBS containing 4% formaldehyde and 2% glutaraldehyde at 4°C for 2 hours. The fixed samples were then gradually dehydrated in a graded ethanol series (30%, 50%, 70%, 95%), with each step performed at 4°C for 15 min. This was followed by three washes in 100% ethanol, each at 4°C for 15 min. The samples were further dehydrated using a Ladd CPD3 Critical Point Dryer and subsequently coated with a thin layer of gold using a Denton Desk V Sputter Coater (Denton Vacuum, USA). Basidia and basidiospores were observed with a scanning electron microscope equipped with an EDS detector (Apreo S, ThermoFisher, USA).

### Nanopore assembly and meiotic recombination analysis

Nanopore sequencing of the reference isolates CSB6039 and CSB6273 was performed using the MinION sequencing device with multiplexing. The DNA for nanopore sequencing was isolated and sequenced as described previously (15, 74). The sequencing was performed with nanopore R10 flow cell with native barcoding kit (SQK-NBD114.24). The sequencing data was processed by basecalling using dorado and genome assembly was performed using Canu v2.0. The assembled genomes were validated for continuity by mapping nanopore reads to the reference assembly using minimap2 as well as by synteny comparison with the already existing reference CBS6039 genome. This resulted in the final genome assembly that consisted of 27 out of 28 telomere repeats with rDNA repeat sequence signatures being present at one end of chromosome 9.

The DNA sequencing reads for the meiotic progeny were quality trimmed using trim_galore (https://www.bioinformatics.babraham.ac.uk/projects/trim_galore/). The trimmed reads for each sample were aligned to the reference de novo CBS6039 genome using bowtie2 (75) with default parameters to obtain sorted BAM files. The BAM files were then used as input for variant calling using GATK (76) in two steps. First, duplicate reads were marked using GATK MarkDuplicates, resulting in DEDUP BAM files. Second, variants were called from these files using GATK HaplotypeCaller and filtered using GATK VariantFilteration to only retain variants with a frequency of 0.9 and coverage of at least 90 reads. The VCF files thus obtained were then plotted against the reference CBS6039 genome in R for visualization and recombination analysis.

### RNA extraction, RNA sequencing, and qRT-PCR assay

To prepare the RNA, cells of tested strains were spotted on V8 (pH=5) agar and incubated for 48 hours same as mating procedures. In parallel, another set was prepared by spotting and incubating cells on YPD agar. Then cells were harvested by using pipet tips and quickly frozen at −80°C until RNA extraction. Three cultures of each strain were grown to provide three biological replicates. Cell disruption was achieved mechanically using Zirconia beads (Ambion, Fisher Scientific, Waltham), and RNA extraction was performed using a 25:24:1 phenol: chloroform: isoamyl alcohol method combined with a Qiagen RNeasy Mini Kit (Qiagen, catalog number 74104). RNA sequencing was performed at Duke University’s Sequencing and Genomic Technology core facility using the Illumina platform. RNA library was prepared by using Watchmaker RNA Library Prep Kits (7BK0002-096),

As for quantitative RT-PCR, a total amount of 1 μg RNA per sample was reverse transcribed to cDNA after DNase I treatment using the Thermofisher Maxima™ H Minus cDNA Synthesis Master Mix (Cat. no M1681). Then, real-time PCR was performed using Power SYBR Green PCR Master Mix (Cat. no 4367659). Relative mRNA levels were normalized to the *ACT1* gene and compared using the threshold cycle ΔΔ*C_T_* method. Differences between cultures were analyzed with the T-test or one-way ANOVA test.

### RNA-sequencing analysis

The sequencing reads were checked for quality and trimmed using trim_galore (https://www.bioinformatics.babraham.ac.uk/projects/trim_galore/). The trimmed and validated reads were mapped to CBS6039 and CBS6273 reference genomes, separately, using the STAR aligner (77) with defined parameters (--outFilterMultimapNmax 10 --alignSJDBoverhangMin 1 --outFilterMismatchNmax 10 --outFilterMismatchNoverReadLmax 0.04 --alignIntronMax 10000 --alignMatesGapMax 1000000). The resulting BAM files were then analyzed for counting reads per gene using “bedtools coverage” (78). The gene count files were used for the DESeq2 analysis in R with the custom scripts, and differentially expressed genes with a log2 fold-change of >1 and p-value of <0.05 were identified. The resulting gene list was split into up-regulated and down-regulated gene sets, which were further analyzed for Gene Ontology enrichment analysis with Uniprot and STRING network analysis.

The promoter sequences for motif analysis were extracted from the reference genome as a 1000-bp sequence upstream of ATG (defined as the start of the first CDS). The extracted sequences were analyzed with Homer “findMotifs.pl” with default parameters. Promoter sequences for all *C. amylolentus* genes (>8000 sequences) were provided as the background set for enrichment analysis. Three separate enrichment analyses were performed for only up-regulated genes, only down-regulated genes, or a combination of both. Identified motifs in each set were scanned for the presence/absence in specific promoters using Homer “scanMotifGenomeWide.pl” feature and custom R scripts.

## Acknowledgements

We are grateful to Anna Floyd Averette for exceptional lab management and technical support, Marco Dias Coelho for technical support with genome sequences and annotations, and current Heitman lab members for their continued interest and many helpful discussions. We thank Dr. Aaron Mitchell from University of Georgia who provided valuable advice for the CRISPR-Cas9 system development.

## Data availability

All sequencing data generated in this study have been deposited in the NCBI database under BioProject accession number PRJNA1331836. The custom R scripts used are available on GitHub (https://github.com/vikasyadavsci/CRAM-RNA-seq-analysis-script)

