## Supplementary figures for "Dissecting the homeodomain *MAT* locus and engineering novel tripolar and bipolar mating systems in *Cryptococcus amylolentus*"

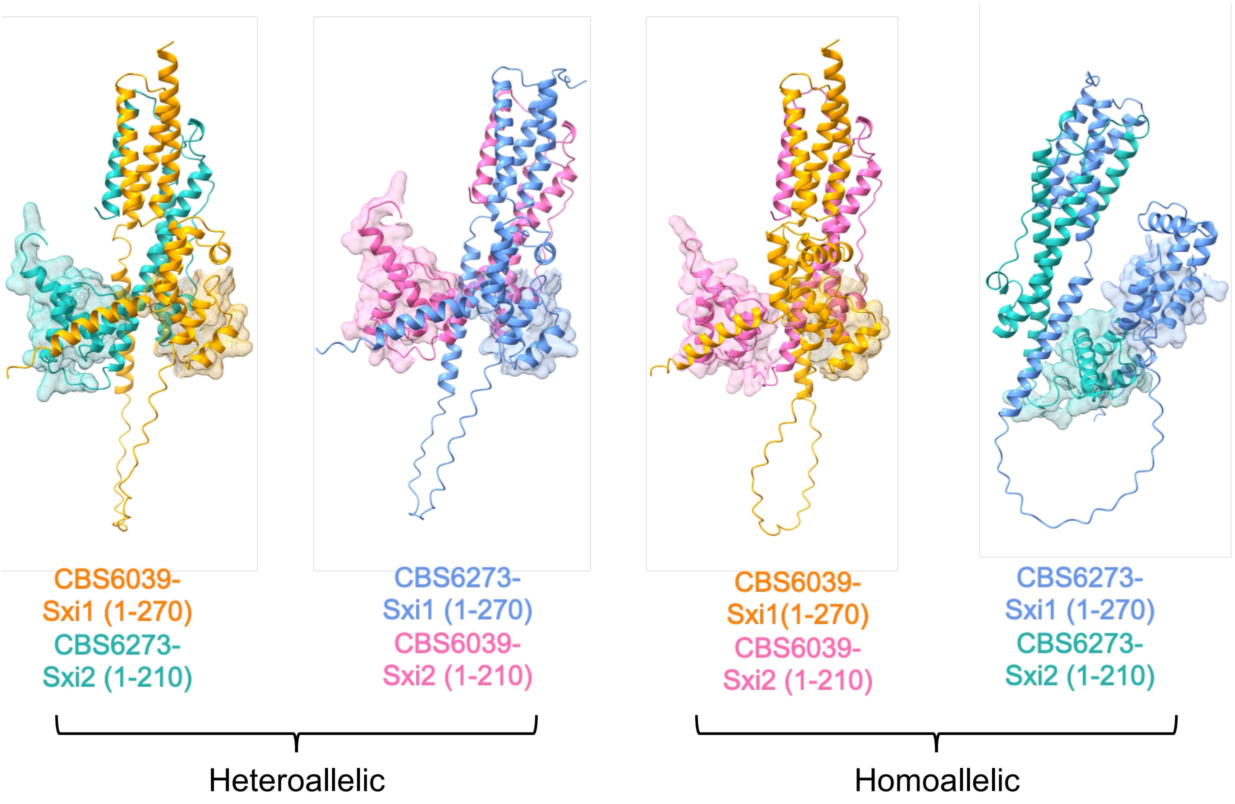


**Figure S1. AlphaFold modeling of *C. amylolentus* Sxi1 and Sxi2 interactions.**

AlphaFold modeling of *C. amylolentus* Sxi1 and Sxi2 interactions was performed based on current annotations. Much ribbon structure is distributed at the peripheral area, the interaction interfaces, mainly between their N-terminal regions, are located at the central area (shown in this figure). AlphaFold modelled structures suggest non-self (heteroallelic) heterodimers and self (homoallelic) heterodimers of the Sxi1 and Sxi2 between and within these two isolates. There was no predicted homodimer formation between Sxi1 or Sxi2 proteins. Predicted homeobox motifs identified in the AlphaFold model are highlighted by shading.

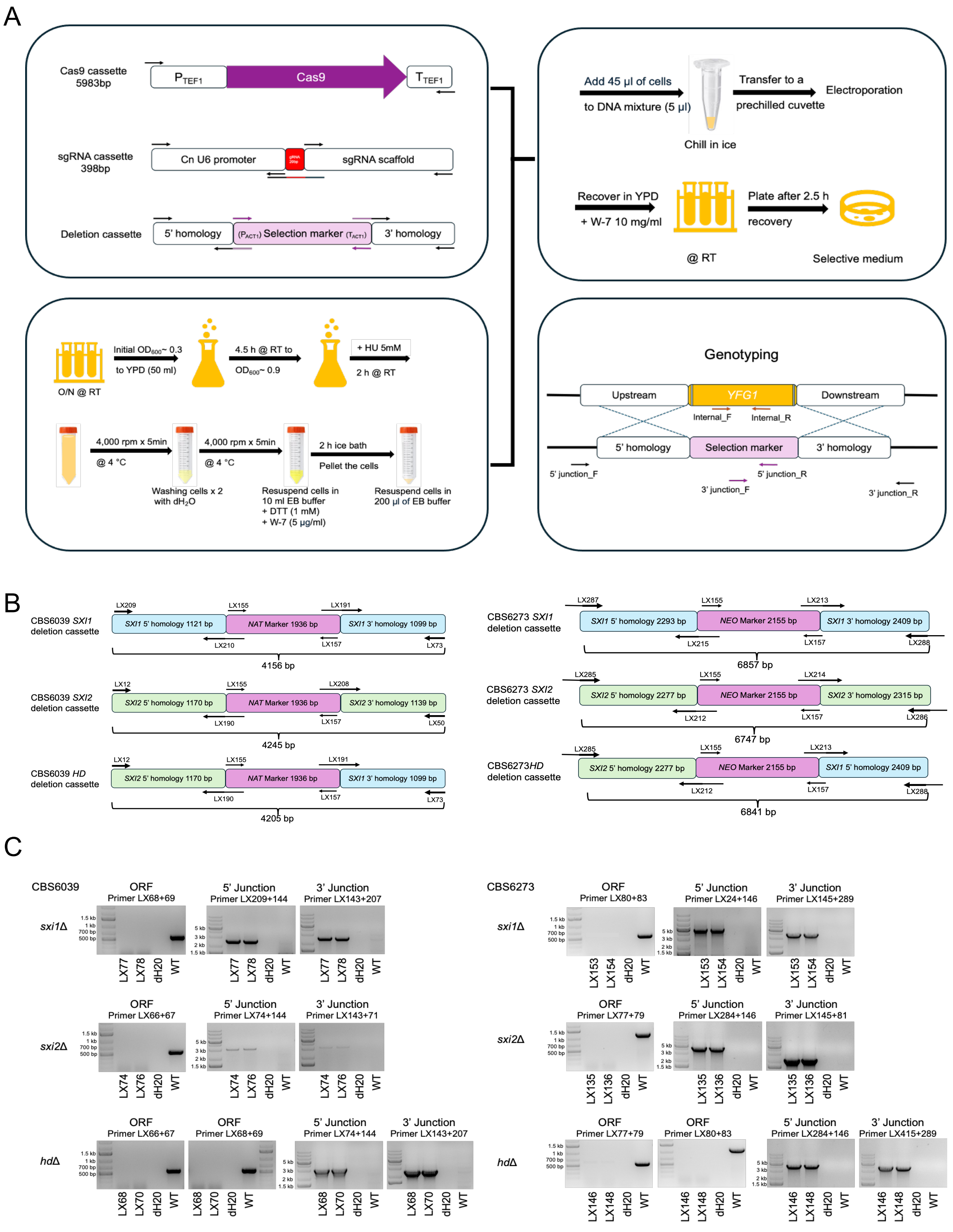

A
Cas9 cassette
5983bp
PTEF1
Cas9
TTEF1
sgRNA cassette
398bp
Cn U6 promoter
sgRNA scaffold
Deletion cassette
5' homology
(PACT1) Selection marker (TACT1)
3' homology
Add 45 µl of cells
to DNA mixture (5 µl)
Chill in ice
Transfer to a
prechilled cuvette
Electroporation
Recover in YPD
+ W-7 10 mg/ml
@ RT
Plate after 2.5 h
recovery
Selective medium
O/N @ RT
Initial OD600~ 0.3
to YPD (50 ml)
4.5 h @ RT to
OD600~ 0.9
+ HU 5mM
2 h @ RT
4,000 rpm x 5min
@ 4 °C
Washing cells x 2
with dH2O
4,000 rpm x 5min
@ 4 °C
Resuspend cells in
10 ml EB buffer
+ DTT (1 mM)
+ W-7 (5 µg/ml)
2 h ice bath
Pellet the cells
Resuspend cells in
200 µl of EB buffer
Genotyping
Upstream
YFG1
Downstream
Internal_F
Internal_R
5' homology
Selection marker
3' homology
5' junction_F
5' junction_R
3' junction_F
3' junction_R
B
CBS6039 SXI1
deletion cassette
LX209
LX155
LX191
SXI1 5' homology 1121 bp
NAT Marker 1936 bp
SXI1 3' homology 1099 bp
LX210
LX157
LX73
4156 bp
CBS6039 SXI2
deletion cassette
LX12
LX155
LX208
SXI2 5' homology 1170 bp
NAT Marker 1936 bp
SXI2 3' homology 1139 bp
LX190
LX157
LX50
4245 bp
CBS6039 HD
deletion cassette
LX12
LX155
LX191
SXI2 5' homology 1170 bp
NAT Marker 1936 bp
SXI1 3' homology 1099 bp
LX190
LX157
LX73
4205 bp
CBS6273 SXI1
deletion cassette
LX287
LX155
LX213
SXI1 5' homology 2293 bp
NEO Marker 2155 bp
SXI1 3' homology 2409 bp
LX215
LX157
LX288
6857 bp
CBS6273 SXI2
deletion cassette
LX285
LX155
LX214
SXI2 5' homology 2277 bp
NEO Marker 2155 bp
SXI2 3' homology 2315 bp
LX212
LX157
LX286
6747 bp
CBS6273HD
deletion cassette
LX285
LX155
LX213
SXI2 5' homology 2277 bp
NEO Marker 2155 bp
SXI1 5' homology 2409 bp
LX212
LX157
LX288
6841 bp
C
CBS6039
sxi1Δ
ORF
Primer LX68+69
5' Junction
Primer LX209+144
3' Junction
Primer LX143+207
LX77 LX78 dH20 WT
sxi2Δ
ORF
Primer LX66+67
5' Junction
Primer LX74+144
3' Junction
Primer LX143+71
LX74 LX76 dH20 WT
hdΔ
ORF
Primer LX66+67
ORF
Primer LX68+69
5' Junction
Primer LX74+144
3' Junction
Primer LX143+207
LX68 LX70 dH20 WT
CBS6273
sxi1Δ
ORF
Primer LX80+83
5' Junction
Primer LX24+146
3' Junction
Primer LX145+289
LX153 LX154 dH20 WT
sxi2Δ
ORF
Primer LX77+79
5' Junction
Primer LX284+146
3' Junction
Primer LX145+81
LX135 LX136 dH20 WT
hdΔ
ORF
Primer LX77+79
ORF
Primer LX80+83
5' Junction
Primer LX284+146
3' Junction
Primer LX415+289
LX146 LX148 dH20 WT
1.5 kb
1 kb
700 bp
500 bp
5 kb
3 kb
2 kb
1.5 kb

**Figure S2. CRISPR-Cas9 system developed for *C. amylolentus*.**

(**A)** Workflow of the Transient CRISPR-Cas9 coupled with Electroporation (TRACE) system developed for *C. amylolentus.* Briefly, the CRISPR-Cas9 system is composed of (1) the Cas9 cassette, which is expressed under the regulation of the native promoter and terminator of the *C. amylolentus TEF1* gene, (2) the sgRNA cassette containing the U6 promoter, gRNA (20 bp), scaffold and 6xT terminator; (3) the repair DNA template of 5' and 3' homology fused to a dominant drug marker adapted with the native promoter and terminator of the *C. amylolentus ACT1* gene. During competent cell preparation, Hydroxyurea (HU), an S-phase inhibitor, and W-7 (hydrochloride), an inhibitor of the NHEJ DNA repair pathway, were included to promote homologous recombination. Transformants were validated by genomic DNA genotyping with specific primer sets targeting the internal region of the gene, 5' junction and 3' junction regions. **(B)** Deletion cassettes of individual *SXI1* or *SXI2* or the combination of both. For mutant construction in CBS6039 (left), fragments of 5' homology, selection marker, and 3' homology were amplified, purified and then fused by junction PCR with the indicated primer sets. For mutant construction in CBS6273 (right), fragments were fused by NEBuilder® HiFi DNA Assembly master mix, then amplified with the resulting plasmid DNA as template. **(C)** Genotyping validation of mutant construction of *sxi1*Δ, *sxi2*Δ and *hd*Δ mutants in both CBS6039 (left) and CBS6273 (right). The Open Reading Frame (ORF) of the *SXI1* or *SXI2* genes, 5' junction and 3' junction regions were detected with the indicated primer sets.

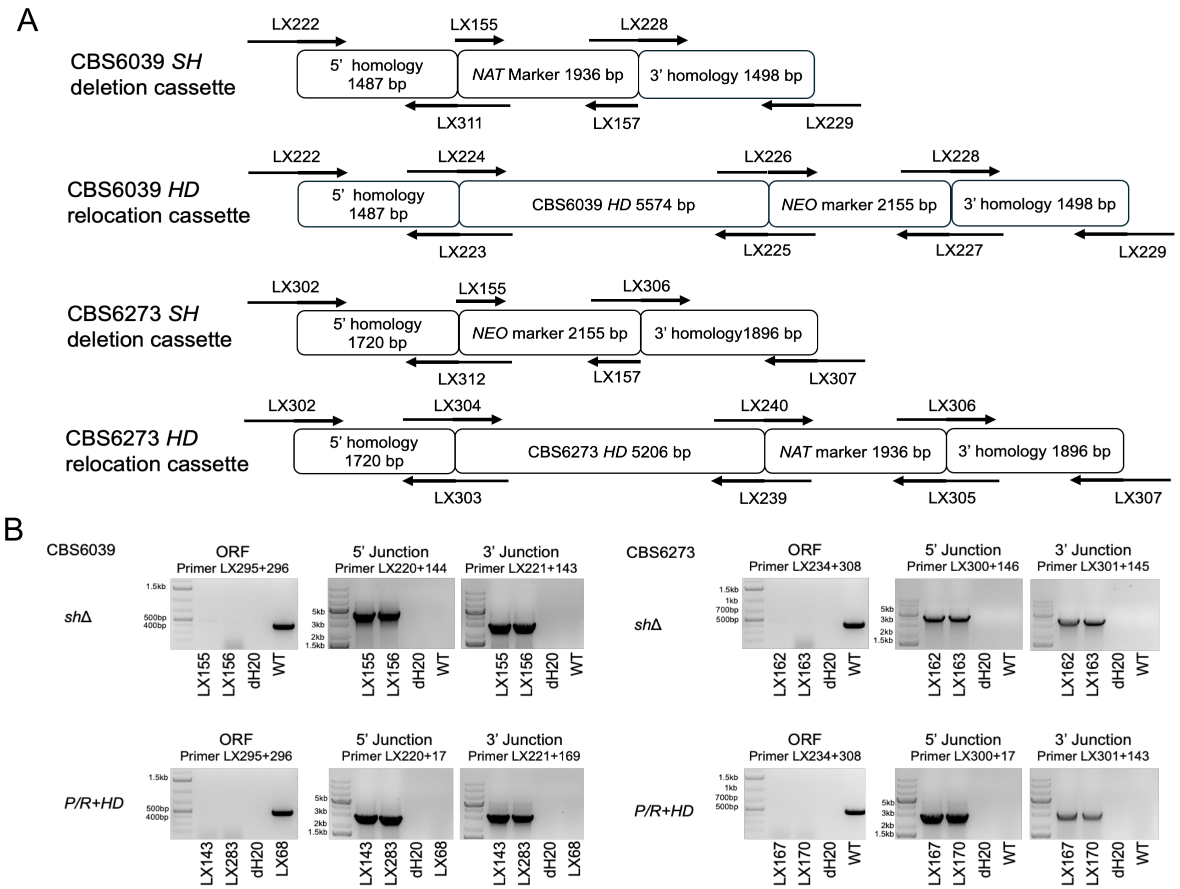


**Figure S3. Construction of *HD* relocated strains in CBS6039 and CBS6273.**

(**A)** Schematic graph (left) of the DNA repair cassettes for strain generation. These cassettes were assembled to a vector pBluescript KS (-) with the NEBuilder® HiFi DNA Assembly master mix. *HD* relocation cassettes (>11 kb) were purified from plasmids with restriction enzymes XbaI and EcoRI. The *SH* deletion cassettes were amplified from plasmid DNA template. (**B)** Genotyping validation of mutant construction of *sh*Δ and *P/R+HD* mutants in both the CBS6039 and the CBS6273 backgrounds. PCRs of *SH* region, 3' junction and 5' junction were conducted with primer sets as indicated.

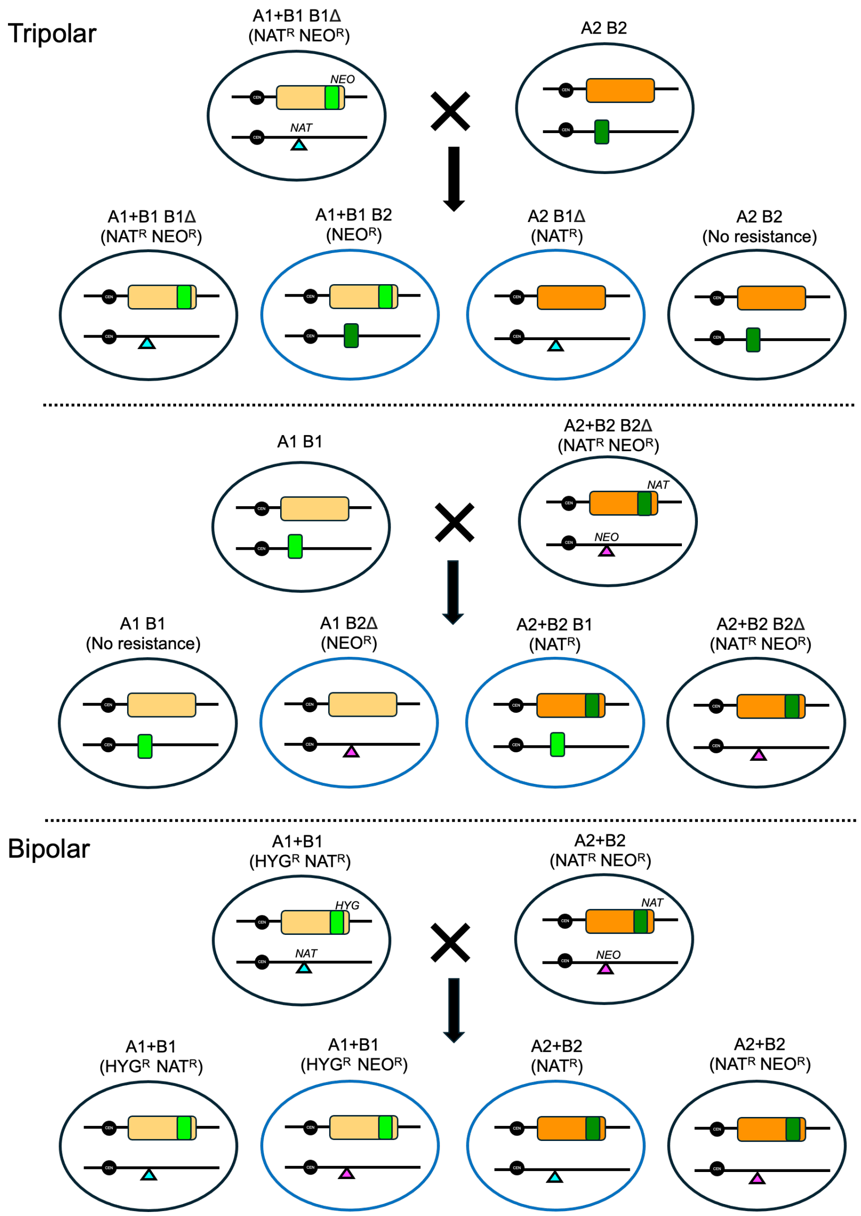


**Figure S4. Diagrams depicting *MAT* loci segregation in tripolar and bipolar crosses.**

Illustrated here are the segregation of the mating types among the progeny in the tripolar and bipolar crosses involving the engineered *C. amylolentus* isolates containing modified *MAT*. In the wild-type strains, the yellow and orange rectangles indicate the A1 and A2 *P/R* alleles, while the light green and dark green rectangles indicate the B1 and

B2 *HD* alleles, respectively. In the engineered "A1+B1 B1Δ" strain, the light blue triangle indicates the deletion of endogenous B1 *HD* genes with the *NAT* marker, and the light green rectangle within the *P/R* locus indicates the insertions of the B1 *HD* genes using the *NEO* selective marker. In the engineered "A2+B2 B2Δ" strain, the purple triangle indicates the deletion of endogenous B2 *HD* genes with the *NEO* marker, and the dark green rectangle within the *P/R* locus indicates the insertions of the B2 *HD* genes using the *NAT* selective marker.

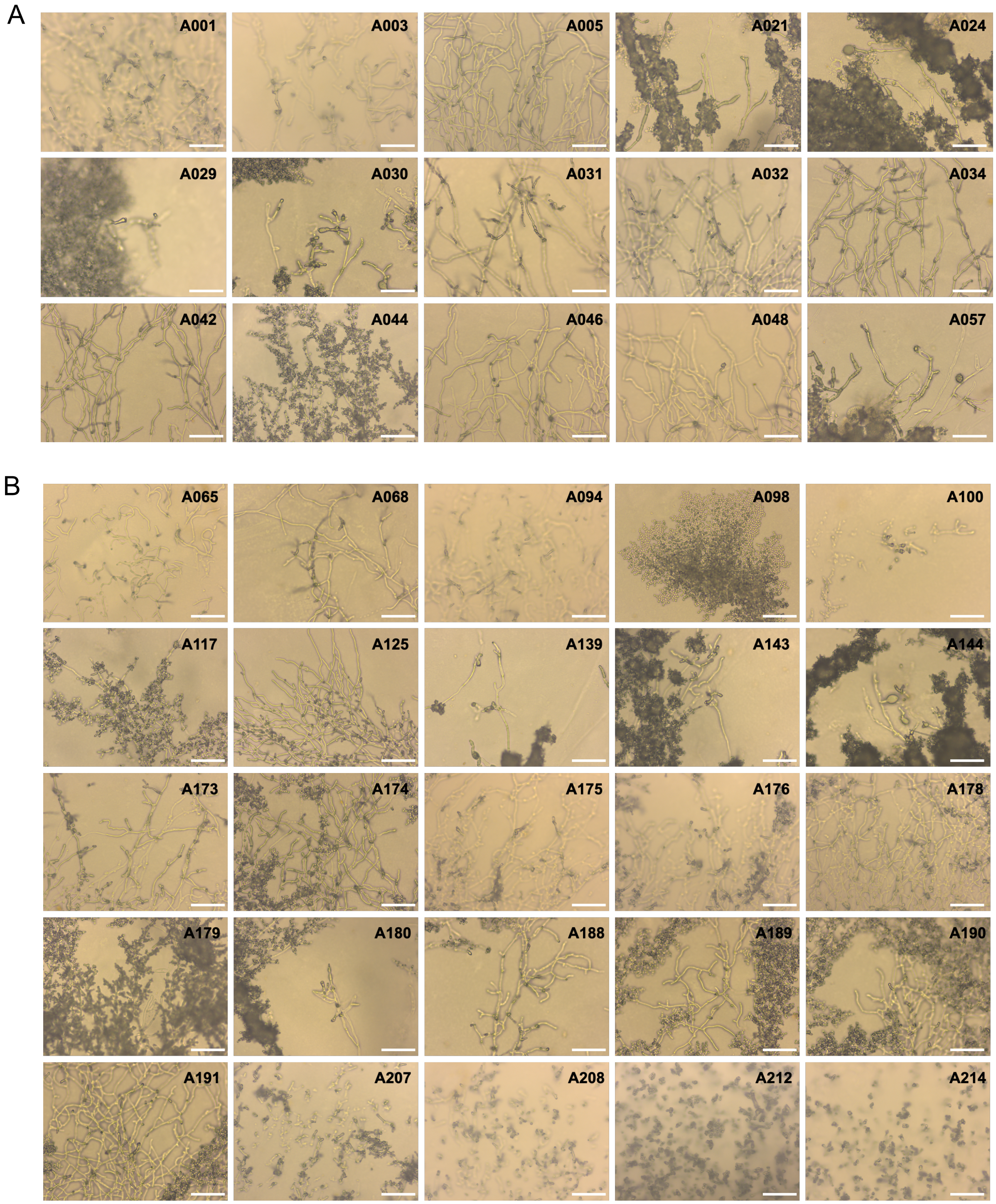


**Figure S5. Phenotypic analysis of F1 progeny harboring two sets of *HD* genes.**

**(A)** Solo culture of 15 F1 progeny inherited the “A2+B2 B1” mating type growing on V8 (pH=5) medium and incubated in the dark at room temperature for 2 weeks. Scale bar:

100 μm. **(B)** Solo culture of 20 F1 progeny inherited the “A1+B1 B2” growing on V8 (pH=5) medium and incubated in the dark at room temperature for 2 weeks. Scale bar: 100 μm.

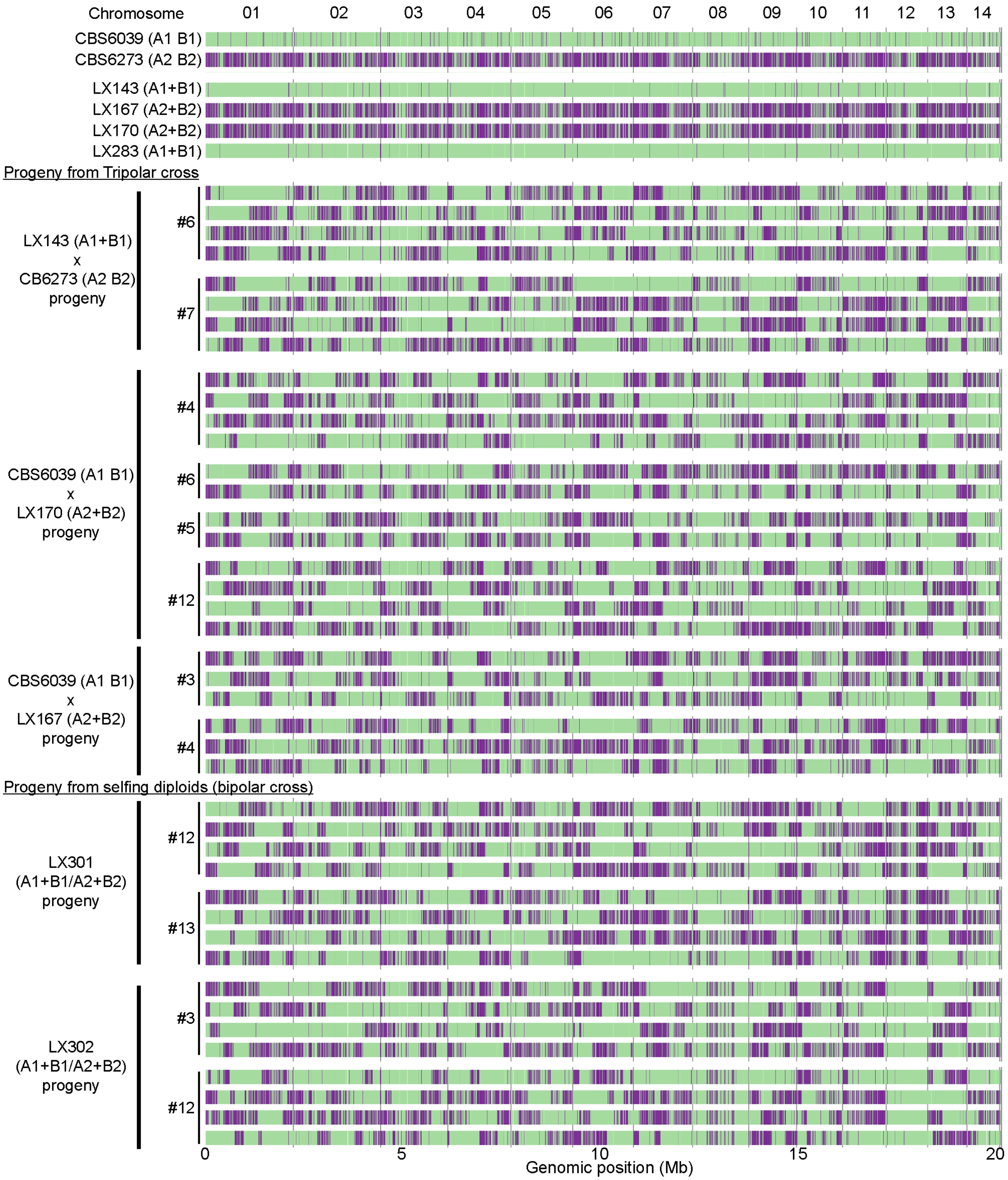


**Figure S6. Variant mapping illustrates genome-wide meiotic recombination in progeny from tripolar and bipolar crosses in *C. amylolentus*.**

SNP patterns of progeny from tripolar and bipolar crosses reveal crossovers consistent with meiotic recombination. Green and purple correspond to variants from CBS6039 and CBS6273, respectively. Parental strains of different mating types and their respective progeny are shown on the left. The numbers of basidia sampled are indicated (#) and correspond to those listed in Tables 1 and 2. For tripolar crosses, both representative basidia and random basidiospores were analyzed; for selfing diploids (A1+B1/A2+B2), eight progeny from two basidia each were examined. *P/R* locus and centromeric (*CEN*) regions are highlighted in shade.

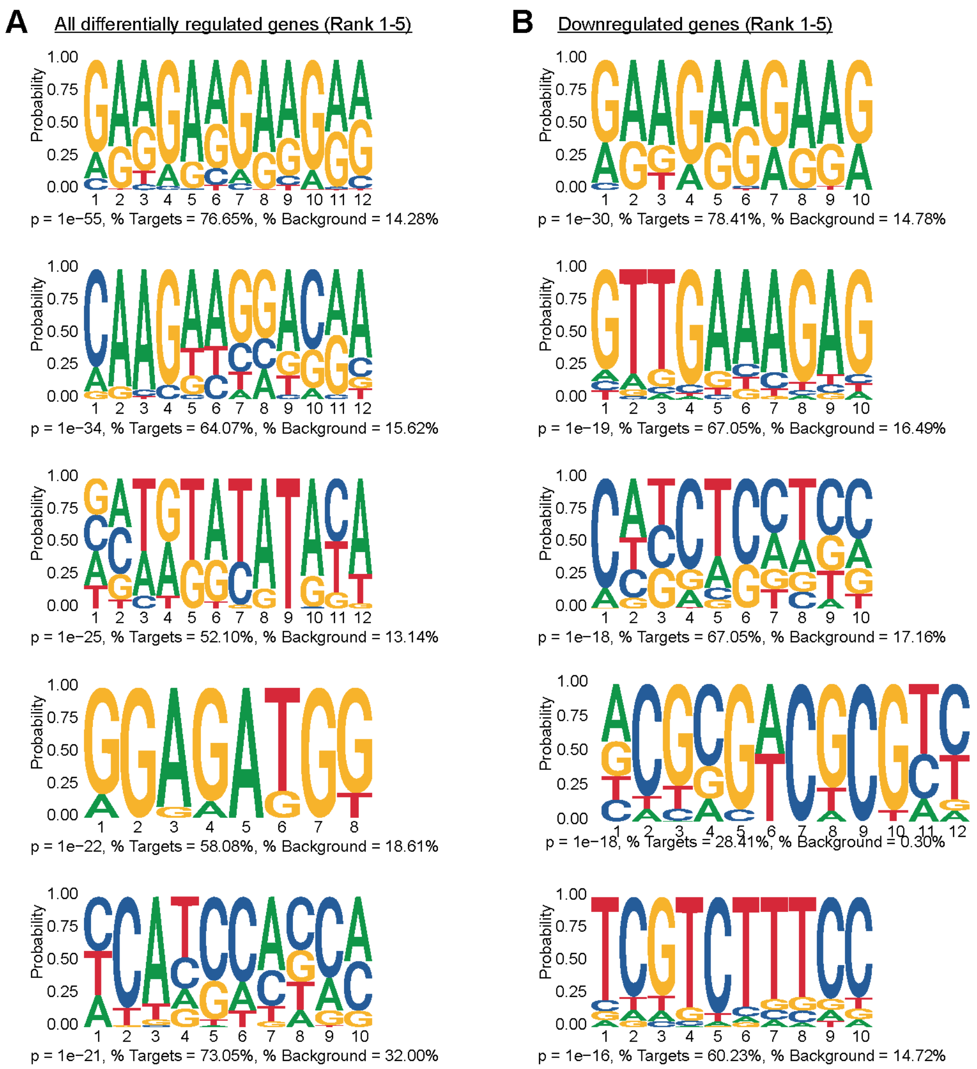


**Figure S7. Predicted binding motifs of Sxi1-Sxi2 responsive genes under mating inducing conditions.**

Motif enrichment analysis was conducted using MEME on the promoter regions (1 kb upstream of the start codon) of all differentially expressed genes **(A)** and down-regulated genes **(B)** in *hd*Δ mutant compared with the wild type under mating inducing conditions. The top 5 enriched sequence motifs (rank 1-5) are shown as sequence logos, representing potential binding sites associated with Sxi1–Sxi2–dependent gene regulation. Motifs are ranked by statistical significance (p value), and the coverage of targets (%) and background (%) are indicated below each motif logo.
